# Tumour Suppressor Parafibromin/Hyrax Governs Cell Polarity and Centrosome Assembly in Neural Stem Cells

**DOI:** 10.1101/2021.07.06.451232

**Authors:** Qiannan Deng, Cheng Wang, Chwee Tat Koe, Jan Peter Heinen, Ye Sing Tan, Song Li, Cayetano Gonzalez, Wing-Kin Sung, Hongyan Wang

**Affiliations:** Neuroscience & Behavioral Disorders Programme, Duke-NUS Medical School, 8 College Road, Singapore 169857; Institute for Research in Biomedicine, The Barcelona Institute of Science and Technology, Baldiri Reixac 10, 08028 Barcelona, Spain; Institució Catalana de Recerca i Estudis Avançats, ICREA, Passeig Lluıs Companys 23, 08010 Barcelona, Spain; Genome Institute of Singapore, 60 Biopolis Street, Genome, Singapore 138672; Department of Computer Science, National University of Singapore, Singapore; Dept. of Physiology, Yong Loo Lin School of Medicine, National University of Singapore, Singapore 117597; NUS Graduate School - Integrative Sciences and Engineering Programme (ISEP), National University of Singapore, Singapore 119077

**Keywords:** *Drosophila*, neural stem cells, neuroblasts, asymmetric division, microtubule growth, centrosome assembly

## Abstract

Neural stem cells (NSCs) divide asymmetrically to balance their self-renewal and differentiation, an imbalance in which can lead to NSC overgrowth and tumour formation. The functions of Parafibromin, a conserved tumour suppressor, in the nervous system are not established. Here, we demonstrate that *Drosophila* Parafibromin/Hyrax (Hyx) inhibits NSC overgrowth by governing cell polarity. Hyx is essential for the apicobasal polarization of NSCs, through its role in the asymmetric distribution of polarity proteins. *hyx* depletion results in the symmetric division of NSCs, leading to the formation of supernumerary NSCs in the larval brain. Importantly, we show that human Parafibromin can fully rescue NSC overgrowth and cell polarity defects in *Drosophila hyx* mutant brains. We have also discovered a novel role for Hyx in regulating the formation of interphase microtubule-organizing center and mitotic spindles in NSCs. Moreover, Hyx is required for the proper localization of a key centrosomal protein, Polo, and the microtubule-binding proteins Msps and D-TACC in dividing NSCs. Furthermore, Hyx directly regulates the *polo* expression *in vitro*. Altogether, our study provides the first evidence that the brain tumour suppressor-like role and polarity establishing functions of Hyx are mediated by its role in regulating microtubule growth and centrosomal assembly. The new paradigm that Parafibromin orchestrates cell polarization by regulating centrosomal assembly may be relevant to future studies on Parafibromin/HRPT2-associated cancers.

## INTRODUCTION

The asymmetric division of stem cells is a fundamental strategy for balancing self-renewal and differentiation in diverse organisms including humans. The *Drosophila* neural stem cells (NSCs), also known as neuroblasts, have emerged as an excellent model for the study of stem cell self-renewal and tumourigenesis (Doe 2008, Januschke and Gonzalez 2008, Wu, Egger et al. 2008, Neumüller and Knoblich 2009, Chang, Wang et al. 2012). During asymmetric division, each NSC generates a self-renewing NSC and a neural progenitor that can produce neurons and glial cells (Doe 2008). Cell polarity is established by the apically localized Par complex, including atypical PKC (aPKC), Bazooka (the *Drosophila* homologue of Par3), and Par6 (Wodarz, Ramrath et al. 1999, Wodarz, Ramrath et al. 2000, Petronczki and Knoblich 2001), as well as the Rho GTPase Cdc42 (Atwood, Chabu et al. 2007). This protein complex displaces the cell fate determinants Prospero, Numb and their adaptor proteins Miranda (Mira) and Partner of Numb to the basal cortex (Ikeshima-Kataoka, Skeath et al. 1997, Shen, Jan et al. 1997, Lu, Rothenberg et al. 1998, Matsuzaki, Ohshiro et al. 1998, Hannaford, Ramat et al. 2018). Another protein complex, includingPins, heterotrimeric G protein subunit Gαi and their regulators, which is linked to the Par proteins by Inscuteable (Insc), is recruited to the apical cortex during mitosis (Kraut, Chia et al. 1996, Schober, Schaefer et al. 1999, Yu, Morin et al. 2000, Schaefer, Petronczki et al. 2001, Yu, Wang et al. 2005, Bowman, Neumüller et al. 2006, Izumi, Ohta et al. 2006). Upon division, apical proteins segregate exclusively into the larger NSC daughter cell to sustain self-renewal, and basal proteins segregate into the smaller progenitor daughter cell to promote neuronal differentiation (Knoblich 2010, Chang, Wang et al. 2012). Such asymmetric protein segregation is facilitated by the orientation of the mitotic spindle along the apicobasal axis (Berdnik and Knoblich 2002, Lee, Andersen et al. 2006, Gonzalez 2007, Wang, Ouyang et al. 2007, Doe 2008, Chabu and Doe 2009, Wang, Chang et al. 2009, Wang, Li et al. 2011). A failure in asymmetric divisions during development may result in cell fate transformation, leading to the formation of ectopic NSCs or the development of brain tumours (Caussinus and Gonzalez 2005, Bello, Reichert et al. 2006, Betschinger, Mechtler et al. 2006, Choksi, Southall et al. 2006, Lee, Andersen et al. 2006, Lee, Robinson et al. 2006, Lee, Wilkinson et al. 2006, Wang, Somers et al. 2006, Wang, Ouyang et al. 2007, Wang, Li et al. 2011).

The dysregulation of a few cell cycle regulators, such as Aurora-A kinase (AurA), Polo kinase (Polo), and Serine/Threonine protein phosphatase 2A (PP2A) results in disruption to NSC asymmetry and microtubule functions, leading to NSC overgrowth and brain tumor formation (Koe and Wang, Terada, Uetake et al. 2003, Wang, Somers et al. 2006, Wang, Ouyang et al. 2007, Chabu and Doe 2009, Krahn, Egger-Adam et al. 2009, Ogawa, Ohta et al. 2009, Wang, Chang et al. 2009, Januschke and Gonzalez 2010, Kelsom and Lu 2012). Moreover, ADP ribosylation factor like-2 (Arl2), a major regulator of microtubule growth, localizes Mini spindles/XMAP215/ch-TOG and Transforming acidic coiled-coil containing (D-TACC) to the centrosomes to regulate microtubule growth and the polarization of NSCs (Chen, Koe et al. 2016).

Human Parafibromin/Cell division cycle 73 (Cdc73)/hyperparathyroidism type 2 (HRPT2) is a tumour suppressor that is linked to several cancers, including parathyroid carcinomas and hyperparathyroidism-jaw tumour syndrome, head and neck squamous cell carcinomas, as well as breast, gastric, colorectal, and lung cancers (Selvarajan, Sii et al. 2008, Zheng, Takahashi et al. 2008, Newey, Bowl et al. 2009, Zhang, Yang et al. 2015). Somatic mutations in parafibromin have been found in 67-100% of sporadic parathyroid carcinomas (Newey, Bowl et al. 2009). Parafibromin is part of a conserved polymerase-associated factor complex that primarily regulates transcriptional events and histone modification (Wei, Yu et al. 1999, Rea, Eisenhaber et al. 2000). Hyrax (Hyx), *Drosophila* Parafibromin, is essential for embryonic and wing development and is known to positively regulate Wnt/Wingless signaling pathway in wing imaginal discs by directly interacting with β-catenin/Armadillo (Mosimann, Hausmann et al. 2006). Human Parafibromin, but not yeast Cdc73, rescues defects in wing development and the embryonic lethality caused by *hyx* loss-of-function alleles (Mosimann, Hausmann et al. 2006), suggesting that Parafibromin functions during development are conserved across metazoans. Interestingly, Parafibromin is expressed in both mouse and human brains, including the cortex, basal ganglia, cerebellum, and the brainstem (Porzionato, Macchi et al. 2006), suggesting that Parafibromin may play a role in central nervous system (CNS) functions. However, the specific functions of Parafibromin in the nervous system are not established. Here, we investigate the role of Parafibromin/Hyx in the asymmetric division of NSCs during *Drosophila* larval brain development.

## RESULTS

### Loss of *hyx* results in NSC overgrowth in the larval central brain

In a clonal screen of a collection of chromosome 3R mutants induced by ethyl methanesulfonate (EMS) (Ly, Tan et al. 2019), we identified two new *hyrax* (*hyx*) alleles - *hyx^HT622^* and *hyx^w12-46^* - which produce NSC overgrowth phenotype in the central brain of *Drosophila* larvae (Figure 1A-B). *hyx*/*CG11990* encodes a highly conserved, 531-amino acid protein that is homologous to mammalian Parafibromin/Cdc73. The *hyx^HT622^* allele contains a 74-bp deletion from nucleotides 728 (immediately after amino acid 242) to 801, which results in a frameshift mutation and, consequently, the generation of a stop codon at amino acid 248. This likely produces a truncated Hyx protein. The other *hyx* allele, *hyx^w12-46^,* carries two point mutations at nucleotides 1331 (T to G) and 1456 (T to A), which causes amino acid substitutions - Leucine (L) to Arginine (R) at amino acid 444 and Cysteine (C) to Serine (S) at amino acid 486, respectively. As *hyx^HT622^* and *hyx^w12-46^* homozygotes are embryonically lethal, we generated MARCM (mosaic analysis with a repressible cell marker) clones (Lee and Luo 1999) to examine the clonal phenotype at larval stages. Hyx protein in the clones was detected using guinea pig anti-Hyx antibodies that we generated against the N-terminal 1-176 amino acids of Hyx. In wild-type control clones, Hyx was predominantly localized to the nuclei of NSCs and their progeny (Figure S1A). In contrast, Hyx was undetectable in 91.3% and 40% of clones generated from *hyx^HT622^* and *hyx^w12-46^* alleles, respectively, and was dramatically reduced in the rest of the clones (Figure S1A). Therefore, these observations indicate that *hyx^HT622^* and *hyx^w12-46^* are two strong loss-of-function alleles.

**Figure 1.**
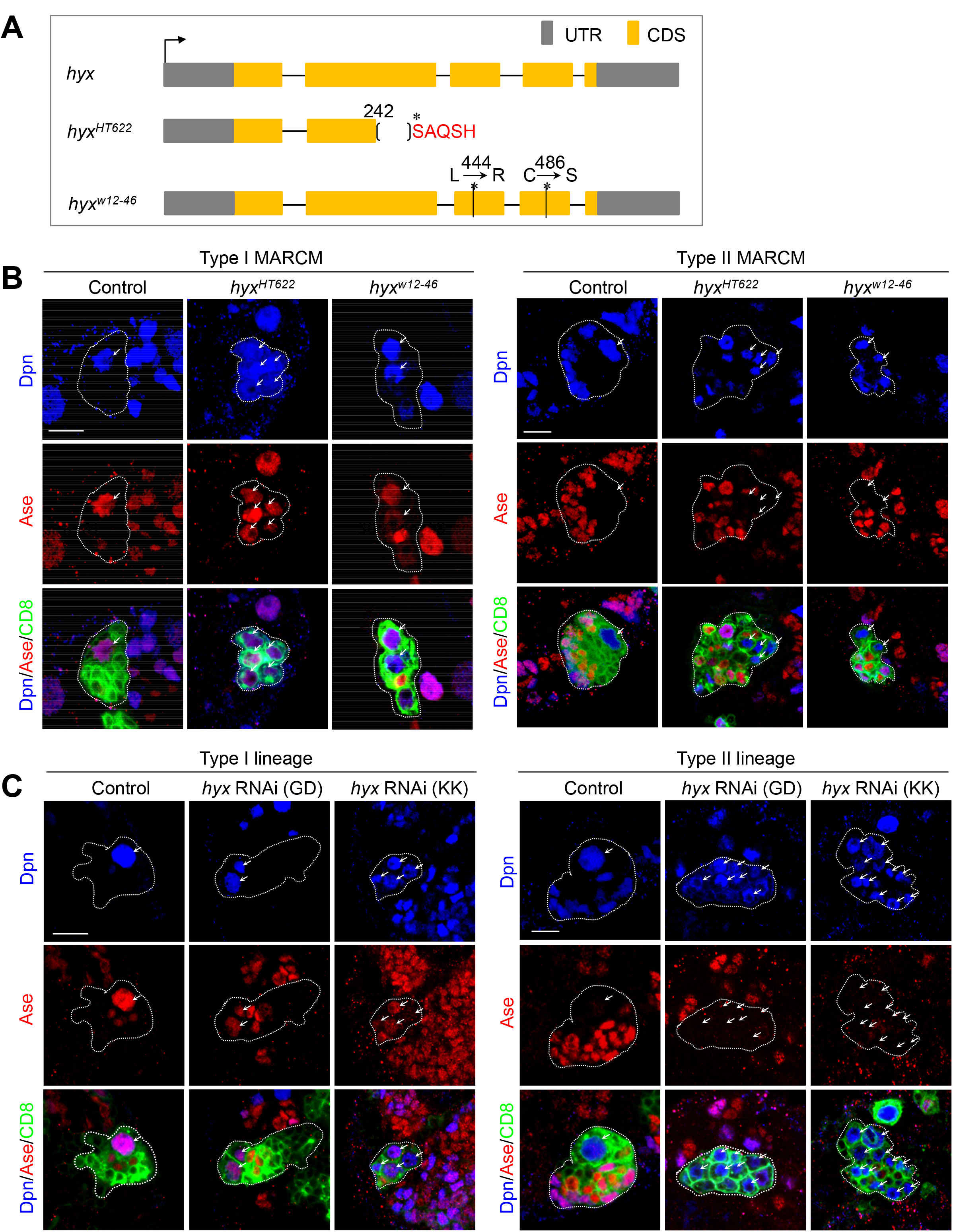
Hyx regulates the homeostasis of central larval brain NSCs. (A) An illustration of two *hyx* EMS alleles. (B) Type I and type II MARCM clones of various genotypes were labeled for Dpn, Ase, and CD8-GFP. Multiple NSCs were observed in *hyx*-depleted type I and type II MARCM clones. Type I clones: control (FRT82B), n=61; *hyx^HT622^*, 87%, n=92; *hyx^w12-46^*, 30.8%, n=68. Type II clones: control (FRT82B), n=57; *hyx^HT622^*, 75%, n=75; 73.3%, *hyx^w12-46^*, n=66. (C) Type I and type II NSC lineages of control (*UAS-β-Gal* RNAi), *hyx* RNAi (GD/V28318), and *hyx* RNAi (KK/V103555) under the control of *insc*-Gal4 driver were labeled for Dpn, Ase, and CD8-GFP. Ectopic NSCs were generated in *hyx* RNAi type I NSC lineages (control, 0%, n=57; GD, 43%, n=92; KK, 27%, n=73) and type II NSC lineages (control, 0%, n=64; GD, 72%, n=83; KK, 89.29%, n=56). MARCM clones in B and NSC lineages in C are outlined by dotted lines. Arrows in B, C indicate NSCs. Scale bars: 5µm.

In the central brain of *Drosophila* larvae, there are at least two types of NSCs, both of which divide asymmetrically (Bello, Izergina et al. 2008, Boone and Doe 2008, Bowman, Rolland et al. 2008). Each type I NSC generates another NSC and a ganglion mother cell (GMC) that gives rise to two neurons, while each type II NSC produces a NSC and a transient amplifying cell (also known as an intermediate neural progenitor or INP) which, in turn, go through a few cycles of asymmetric divisions to produce GMCs (Bello, Izergina et al. 2008, Boone and Doe 2008, Bowman, Rolland et al. 2008, Weng, Golden et al. 2010). In the wild-type control, only one NSC is maintained in each of type I or type II MARCM clones (Figure 1B). However, ectopic NSCs were observed in 87% of type I clones and 75% of type II clones generated from the *hyx^HT622^* allele (Figure 1B). Similarly, supernumerary NSCs were observed in both type I and type II NSC lineages from *hyx^w12-46^* clones (Figure 1B). In addition, knockdown of *hyx* by two independent RNAi lines, under the control of a NSC driver *insc*-Gal4, led to the formation of multiple NSCs in both type I and type II lineages (Figure 1C). Moreover, the NSC overgrowth phenotype in *hyx^HT622^* and *hyx^w12-46^* mutants was fully rescued by the overexpression of a wild-type *hyx* transgene (Figure S1B-C). Therefore, our finding shows that *hyx* prevents NSC overgrowth in both type I and type II lineages.

Remarkably, the ectopic NSC phenotype observed in *hyx^HT622^* larval brains was also completely rescued by the overexpression of Parafibromin/HRPT2, the human counterpart of Hyx (Figure S1D). Likewise, the NSC overgrowth phenotype observed in *hyx* knockdown brains was fully restored by the introduction of human Parafibromin/HRPT2 in both type I and type II NSC lineages (Figure S1E). Taken together, Parafibromin/Hyx appears to have a conserved function in suppressing NSC overgrowth.

Parafibromin/Cdc73 (Hyrax/Hyx in *Drosophila*) is a component of the PAF1 complex, an evolutionarily conserved protein complex that functions in gene regulation and epigenetics (Costa and Arndt 2000, Mueller, Porter et al. 2004, Penheiter, Washburn et al. 2005, Van Oss, Cucinotta et al. 2017). The PAF1 complex also consists of other core subunits Paf1 (Antimeros/atms in *Drosophila*), Leo1 (Another transcription unit/Atu in *Drosophila*), Cln three requiring 9 (Ctr9 in *Drosophila*), and Rtf1 (Mueller and Jaehning 2002, Zhu, Mandal et al. 2005). We sought to analyze the function of other components of the PAF1 complex in the larval central brains. Surprisingly, although Ctr9 is required to terminate the proliferation of *Drosophila* embryonic NSCs (Choksi, Southall et al. 2006, Bahrampour and Thor 2016), no ectopic NSCs were observed in *ctr9^12P023^* type I and type II MARCM clones (Figure S1F). Similarly, knockdown of *ctr9, atms*, or *atu* under the control of *insc*-Gal4 did not generate supernumerary NSCs in either type I or type II lineages in the larval central brains (n=5 for all). Interestingly, knocking down *rtf* resulted in weak type II NSC overgrowth phenotype, without affecting type I NSC lineage development (*rtf1* RNAi/BDSC#34586: type II, 31.2%, n=64 and *rtf1* RNAi/BDSC#34850: type II, 7%, n=43). Therefore, our results suggest that Parafibromin/Hyx is a key component of Paf1 complex that prevents NSC overgrowth during *Drosophila* brain development.

### Hyx is essential for the asymmetric division of NSCs

The generation of ectopic NSCs in the absence of Hyx function was not due to INP dedifferentiation, as no type II NSCs were generated in both control and *hyx* RNAi/V103555 derived INP clones (Figure S2A). Next, we assessed whether *hyx* is required for the asymmetric division of NSCs. In wild-type control metaphase NSCs, apical proteins such as aPKC, Insc, Baz/Par3, Par6, and Pins were localized asymmetrically in the apical cortex (Figure 2A-F). By contrast, aPKC in *hyx^HT622^* and *hyx^w12-46^* metaphase NSCs was completely delocalized from the apical cortex to the cytoplasm (Figure 2A-B). Similarly, other apical proteins including Insc, Baz, Par6, and Pins in *hyx^HT622^* metaphase NSCs were no longer localized asymmetrically in the apical cortex, exhibiting weak or punctate signals in the cytoplasm (Figure 2C-F).

**Figure 2.**
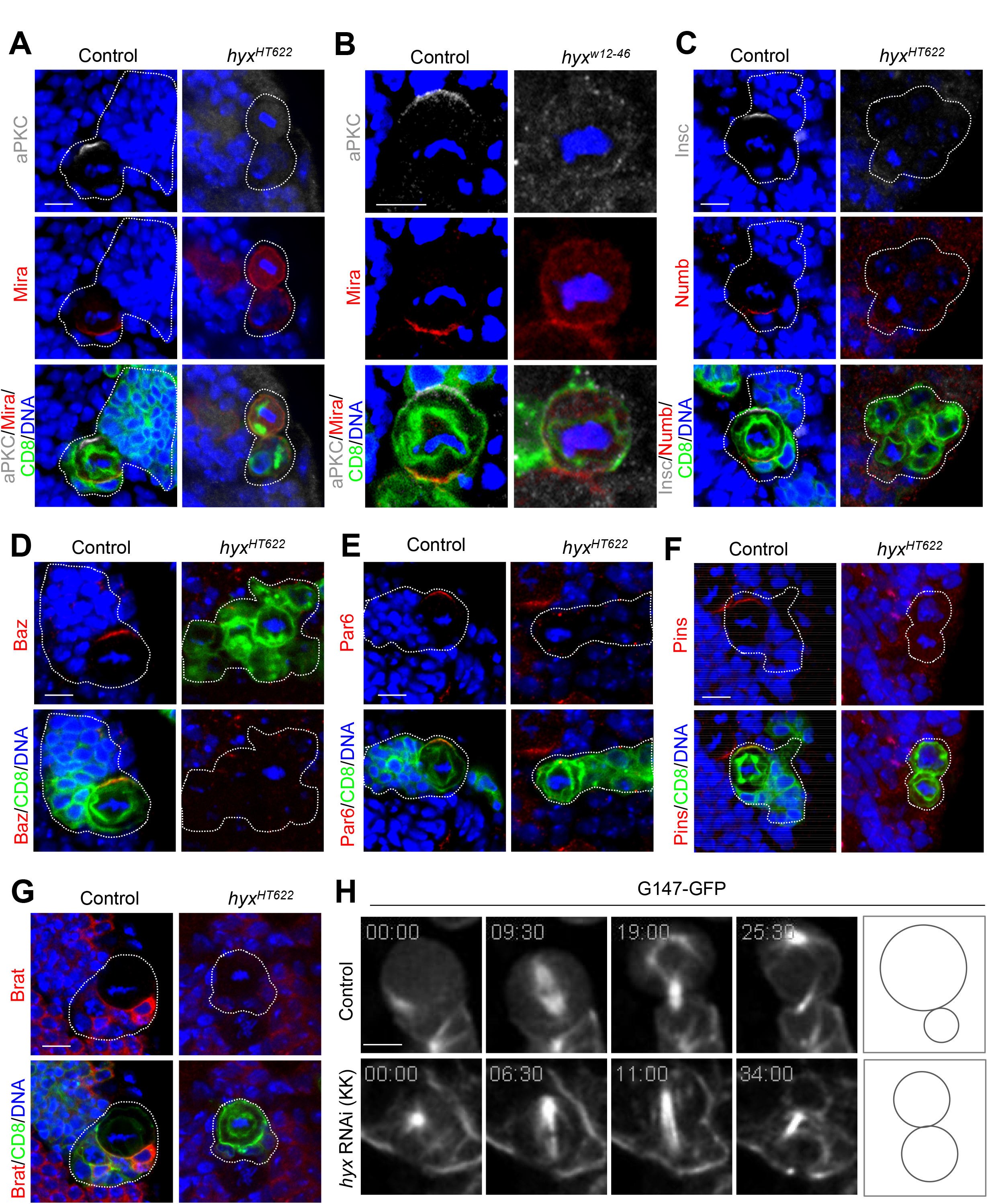
Hyx is essential for asymmetric cell division of NSCs. (A) Metaphase NSCs of control (FRT82B; n=50) and *hyx^HT622^* MARCM clones were labeled for aPKC, Mira, and DNA. *hyx^HT622^*, aPKC delocalization, 100%; Mira delocalization, 81.6%; n=49 for both. (B) Metaphase NSCs of control (FRT82B; n=45) and *hyx^W12-46^* MARCM clones were labeled for aPKC, Mira, and DNA. 100% delocalization of aPKC and Mira in *hyx^W12-46^*, n=50 for both. (C) Metaphase NSCs of control (FRT82B; n=47) and *hyx^HT622^* MARCM clones were labeled for Insc, Numb, and DNA. 100% delocalization of Insc and Numb in *hyx^HT622^*, n=49 for both. (D) Control (FRT82B; n=46) and *hyx^HT622^* metaphase NSCs were labeled for Baz and DNA. 100% Baz delocalization in *hyx^HT622^* (n=49). (E) Control (FRT82B; n=50) and *hyx^HT622^* metaphase NSCs were labelled for Par6 and DNA. 100% delocalization of Par6 in *hyx^HT622^* (n=49). (F) Metaphase NSCs from control (FRT82B) and *hyx^HT622^* were labelled for Pins and DNA. 100% delocalization of Pins in *hyx^HT622^*; n=49 for both genotypes. Clones were marked by CD8-GFP. (G) Metaphase NSCs from control (FRT82B; n=48) and *hyx^HT622^* were labelled for Brat and DNA. 100% Brat delocalization in *hyx^HT622^* (n=50). (H) Still images of time-lapse imaging of control (G147-GFP/+; n=15 and Video S1) and *hyx* RNAi (KK/V103555; n=5 and Video S2) NSCs expressing G147-GFP under the control of *insc*-Gal4 at 48h ALH. MARCM clones are marked by dotted lines (A, C-G). Scale bar: 5 μm.

Next, we examined the localization of basal proteins in *hyx* mutant NSCs. Mira was asymmetrically localized in the basal cortex in 100% of wild-type NSCs during metaphase, but its basal localization was severely disrupted in metaphase NSCs of *hyx^HT622^* and *hyx^w12-46^* clones (Figure 2A-B). Similarly, Numb and Brat lost their asymmetric basal localization, and were observed in the cytoplasm in metaphase NSCs of *hyx^HT622^* clones (Figure 2C, G).

Similarly, *hyx* knockdown by two independent RNAi lines disrupted NSC apicobasal polarity. aPKC, Par6, Baz, Insc, and Pins were delocalized from the apical cortex in all metaphase NSCs of both *hyx* RNAi lines (Figure S2B-G). In addition, the localization of Mira, Numb, and Brat proteins at the basal cortex was also disrupted in metaphase NSCs upon *hyx* knockdown (Figure S2B-C, S2D, S2H). Taken together, our data indicate that Hyx orchestrates NSC polarity by regulating the proper localization of both apical and basal proteins.

Given that polarization of NSCs is an essential prerequisite for their asymmetric division, we sought to investigate whether *hyx* depletion could result in the symmetric division of NSCs. To this end, we took advantage of a microtubule-binding protein Jupiter-GFP (also known as G147-GFP) which is controlled under its endogenous promoter (Morin, Daneman et al. 2001). In live, whole-mount larval brains that expressed G147-GFP, control NSCs always divided asymmetrically to produce two daughter cells with distinct cell sizes (Figure 2H, movie S1). By contrast, all NSCs in *hyx* RNAi divided symmetrically to generate two daughter cells with similar cell sizes (Figure 2H, movie S2). These observations indicate that *hyx*-depleted NSCs divide symmetrically, leading to NSC overgrowth.

### Hyx maintains interphase microtubule asters and the mitotic spindle

When analyzing the asymmetric division, we noticed that *hyx*-depleted NSCs formed shorter mitotic spindles. The mean spindle length in control metaphase NSCs was 9.81 ± 0.94 µm, whereas in *hyx* RNAi (V103555) NSCs, the mean spindle length was dramatically shortened to 6.84 ± 1.46 µm (Figure 3A-B). Consistent with this observation, spindle lengths in metaphase *hyx^HT622^* and *hyx^w12-46^* NSCs were much shorter than that in the control (Figure 3C-D).

**Figure 3.**
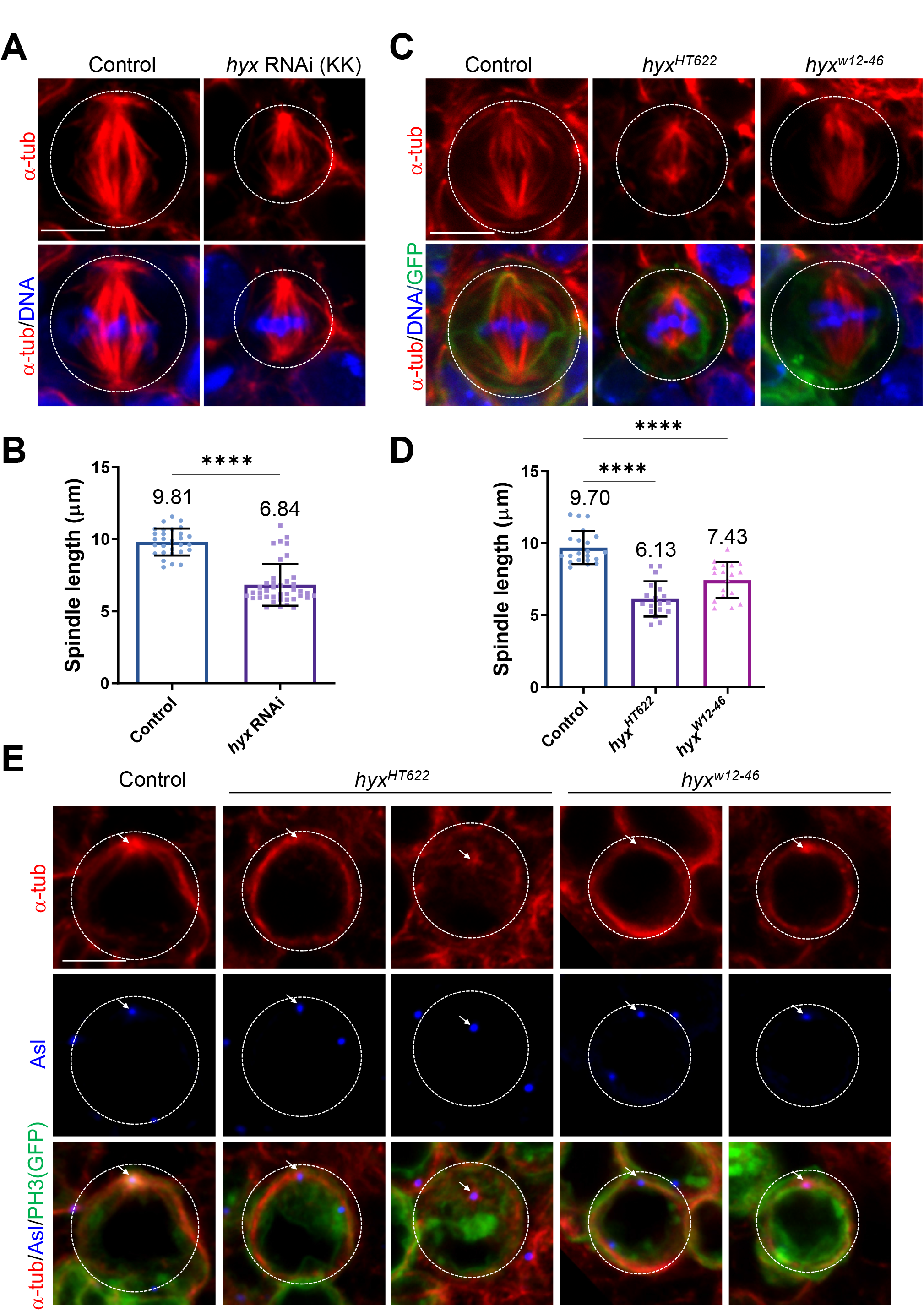
Hyx is required for the formation of the microtubule aster and the mitotic spindle in NSCs. (A) Metaphase NSCs of control (*UAS-β-Gal* RNAi) and *hyx* RNAi (KK/V103555) under the control of *insc*-Gal4 were labelled for α-tub and DNA. (B) Quantification of spindle length (with SD) observed in A. Spindle length: control, 9.81 ± 0.94 µm, n=29; *hyx* RNAi, 6.84 ± 1.46 µm, n=40. (C) NSCs of control (FRT82B), *hyx^HT622^*, and *hyx^W12-46^* MARCM clones were labelled for α-tub and DNA and CD8-GFP. (D) Quantification of spindle length (with SD) observed in C. Spindle length: control, 9.70± 1.13 µm, n=21; *hyx^HT622^*, 6.13 ± 1.22 µm, n=18; *hyx^W12-46^*, 7.43 ± 1.25 µm, n=18. (E) Interphase NSCs from control (FRT82B; n=11), *hyx^HT622^* (n=24), and *hyx^w12-46^* (n=29) MARCM clones were labelled for α-tub, Asl, GFP, and PH3. NSCs failed to form a microtubule aster in 79.2% of *hyx^HT622^* and 34.5% of *hyx^w12-46^* clones. The rest of them formed weak microtubule asters. Cell outlines are indicated by white dotted lines. Arrows indicate the centrosomes. Scale bars: 5 μm.

These observations prompted us to examine whether Hyx is important for microtubule assembly in NSCs. We sought to determine whether Hyx regulates the formation of microtubule asters in interphase NSCs. A wild-type interphase NSC forms one major microtubule aster marked by α-tubulin (α-tub). Asters are assembled by the microtubule-organizing center (MTOC), also known as centrosomes, of cycling NSCs labeled by a centriolar protein called Asterless (Asl; Figure 3E). Strikingly, the vast majority of interphase NSCs in *hyx^HT622^* and *hyx^w12-46^* clones either failed to organize a microtubule aster or formed weak microtubule asters (Figure 3E). Likewise, in *hyx* knockdown clones, 50% of NSCs failed to form microtubule asters and 40.9% of NSCs only assembled weak microtubule asters during interphase (Figure S3A). Overall, we show that Hyx is important for the formation of interphase microtubule asters.

The shortened mitotic spindles and defects in the assembly of microtubule asters upon *hyx* depletion suggested that Hyx might regulate microtubule growth in dividing NSCs. To this end, we performed a microtubule regrowth assay by “cold” treatment of larval brains on ice - for efficiently depolymerizing microtubules in NSCs - followed by their recovery at 25°C to allow microtubule regrowth in the course of time. In both control and *hyx* RNAi interphase NSCs treated with ice (t=0), no astral microtubules were observed and only weak residual microtubules labelled by α-tub remained at the centrosome (Figure 4A). The centrosomes in 76.3% of these *hyx* RNAi interphase NSCs were absent or much smaller in size, suggesting that the MTOC was compromised upon *hyx* depletion (Figure 4A). In control interphase NSCs, robust astral microtubules were observed around the centrosome, at various time points following recovery at 25°C (Figure 4A). By contrast, the vast majority of *hyx* RNAi interphase NSCs reassembled scarce microtubule bundles without detectable MTOCs, even 120 seconds after recovery at 25°C (Figure 4A).

**Figure 4.**
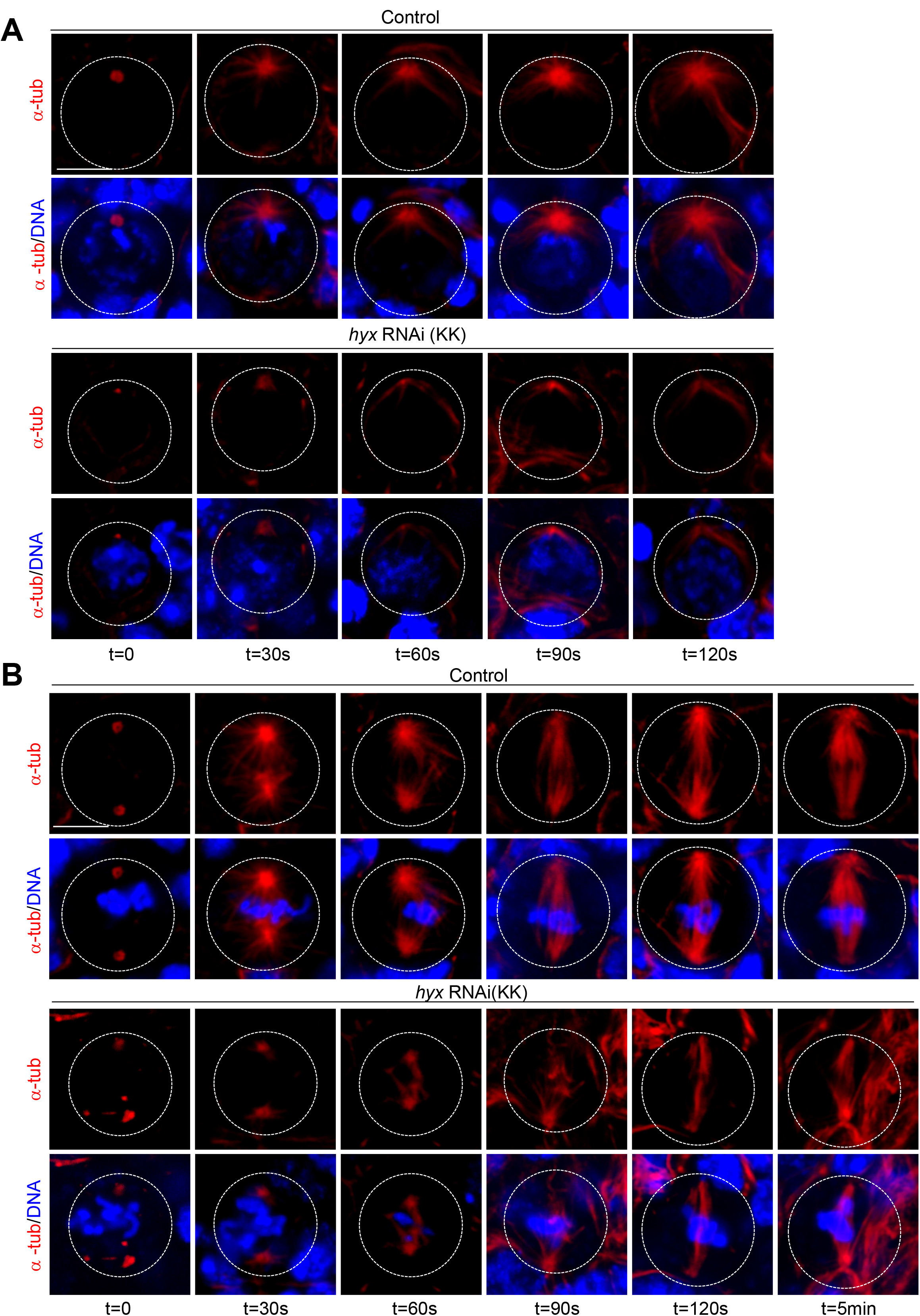
Hyx promotes microtubule growth in NSCs. (A) Control (*UAS-β-Gal* RNAi) and *hyx* RNAi (KK/V103555) interphase NSCs were labelled for α-tub and DNA at various time points after recovery from treatment on ice in microtubule regrowth assay. At 0 second (t=0; ice treatment), no astral microtubules around the centrosome/MTOC were observed in all control (n=74) and *hyx* RNAi NSCs (n=63). At all subsequent time points following the recovery, all control NSCs assembled robust astral microtubules (t=30s, n=31; t=60s, n=11; t=90s, n=27; t=120s, n=26). Astral microtubules were lost in *hyx* RNAi: t=30 s, 71.9%, n=43; t=60 s, 85.7%, n=7; t=90 s, 91.2%, n=46; t=2 min, 86.3%, n=63. (B) Control (*UAS-β-Gal* RNAi) and *hyx* RNAi (KK/V103555) metaphase NSCs were labelled for α-tub and DNA at various time points after recovery from treatment on ice in microtubule regrowth assay. Control: t=0, n=52; t=30 s, n=19; t=60 s, n=9; t=90 s, n=41; *hyx* RNAi, t=0, n=28. Scarce spindle microtubule mass assembled in *hyx* RNAi: t=30 s, 91.7%, n=18; t=60 s, 66.7%, n=3; t=90 s, 69.7%, n=19; t=2 min, 85.4%, n=11. RNAi was controlled by *insc*-Gal4. Cell outlines are indicated by the white-dotted lines. Scale bars: 5 μm.

Next, we examined microtubule regrowth in mitotic NSCs. Upon treatment with ice (t=0), spindle microtubules were effectively depolymerized with residual microtubules marking the centrosomes/spindle poles, in all control and *hyx* RNAi metaphase NSCs (Figure 4B). Consistent with poor centrosome assembly during interphase, the centrosomes of 98.0% of *hyx* RNAi metaphase NSCs were deformed with irregular shapes (Figure 4B). In all control metaphase NSCs, intense spindle microtubules were reassembled around centrosomes and chromosome mass from as early as 30s following recovery at 25°C; the mitotic spindle completely re-formed at 2 min’s following recovery (Figure 4B). In contrast, the majority of *hyx* RNAi metaphase NSCs assembled scarce spindle microtubule mass following recovery; at 2 mins after recovery, only 14.6% of metaphase NSCs formed mitotic spindles, which were still shorter and thinner than the spindles formed in the control NSCs. MTOCs remained weak or absent in these *hyx*-depleted NSCs (Figure 4B). Taken together, we propose that Hyx plays a central role in the formation of interphase microtubule asters and the mitotic spindle by promoting microtubule growth in NSCs.

### Hyx is required for centrosome assembly in dividing NSCs

Since we had demonstrated that Hyx promotes microtubule growth and interphase aster formation in NSCs, we wondered whether Hyx regulates the assembly of the centrosomes, the major MTOC in dividing cells. Each centrosome is composed of a pair of centrioles surrounded by pericentriolar material (PCM) proteins. Centriolar proteins Spindle assembly abnormal 4 (Sas-4), Anastral spindle 2 (Ana2), and Asl are essential for centriole biogenesis and assembly (Varmark, Llamazares et al. 2007, Dzhindzhev, Yu et al. 2010, Wang, Li et al. 2011, Novak, Wainman et al. 2016, Gartenmann, Vicente et al. 2020). In control interphase and metaphase NSCs, Sas-4 was always observed at the centrosomes overlapping with Asl (Figure S3B-C). In response to *hyx* knockdown, the localization of Sas-4 and Asl seemed unaffected at the centrosomes in both interphase and metaphase NSCs (Figure S3B-C).

Wild-type NSCs typically contained two Asl-positive centrioles (Figure S3D-E), as the centrioles are duplicated early in the cell cycle. Strikingly, multiple centrioles marked by Asl were observed at the centrosomes in 47.1% of *hyx^HT622^* clones (Figure S3D-E), 10.0% of *hyx^w12-46^* clones (n=8), and 12.9% of *hyx* RNAi NSCs (Figure S3F). Consistent with these observations, multiple centrioles were detected in 52.0% of S2 cells upon dsRNA treatment against *hyx* (Figure S3G-H), which was significantly higher than what was observed in control cells (Figure S3G-H; 25.1%). These data suggest that Hyx might also affect centriole duplication.

Major PCM proteins γ-tubulin (γ-tub) and centrosomin (CNN, a CDK5RAP2 homolog), are essential for microtubule nucleation and anchoring. γ-tub is a component of the γ-tubulin ring complex (γ-TURC), which is the major microtubule nucleator in dividing cells, including NSCs (Moritz, Braunfeld et al. 1995, Sunkel, Gomes et al. 1995). In control interphase and metaphase NSCs, robust γ-tub was detected at the centrosomes and was observed to co-localize with Asl (Figure 5A). By contrast, γ-tub was absent or significantly reduced at the centrosomes in 95.7% of *hyx^HT622^* and 54.8% of *hyx^w12-46^* interphase NSCs (Figure 5A); the fluorescence intensity of γ-tub was dramatically decreased in these NSCs (Figure 5A, C). In addition, γ-tub was strongly diminished at the centrosomes in 93.1% of *hyx^HT622^* and 70.8% of *hyx^w12-46^* metaphase NSCs (Figure 5B). The fluorescence intensity of γ-tub protein dropped to 0.17-fold in *hyx^HT622^* and 0.53-fold in *hyx^w12-46^* NSCs, respectively (Figure 5B-C). Likewise, γ-tub was strongly reduced at the centrosomes in 90.5% of interphase and 81.3% of metaphase NSCs upon *hyx* RNAi knockdown (Figure S4A-B); moreover, the fluorescence intensity of γ-tub was dramatically decreased at the centrosomes in these NSCs (Figure S4A-C).

**Figure 5.**
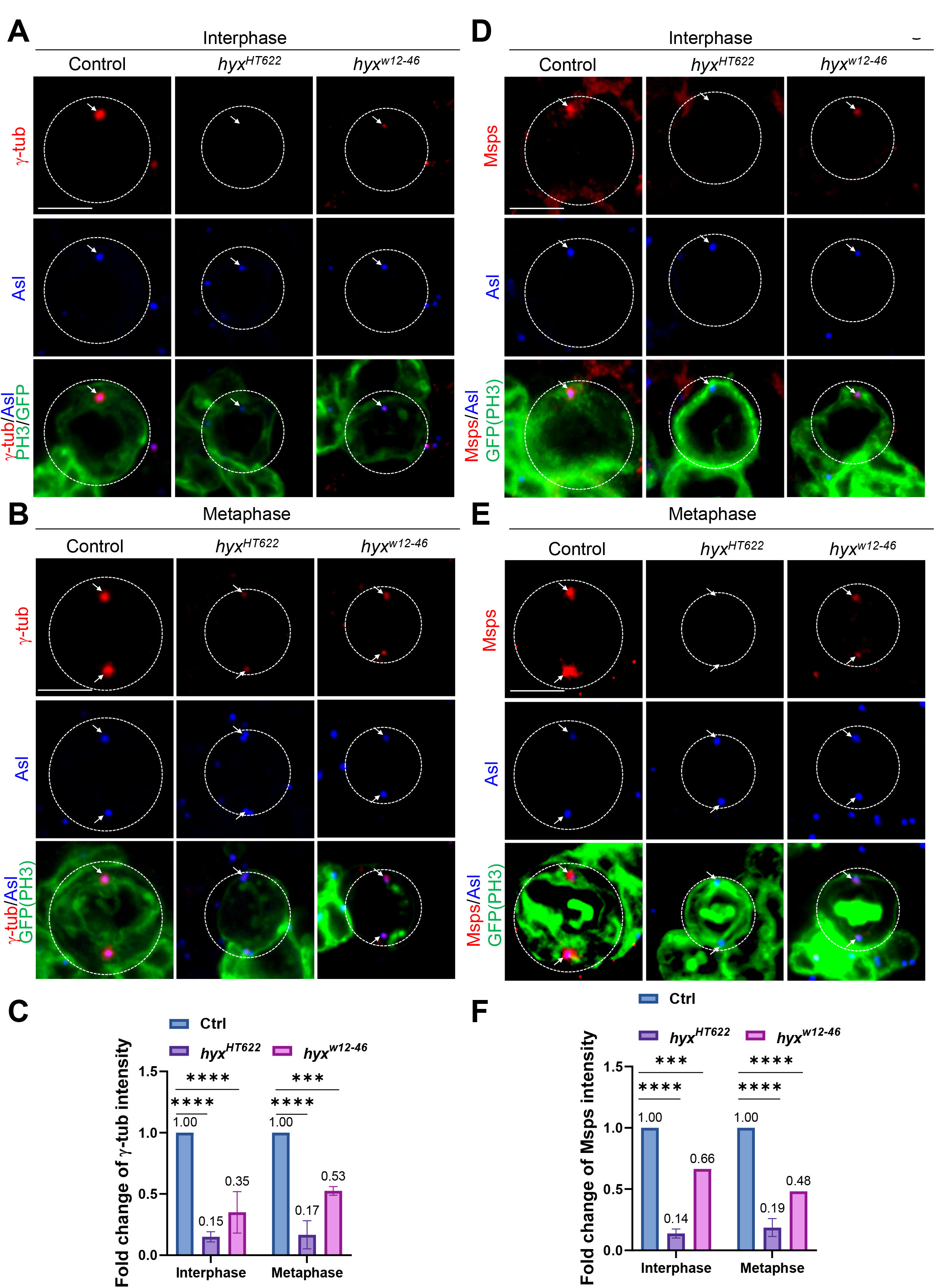
Hyx regulates the localization of centrosomal proteins γ-tubulin and Msps in NSCs. (A) Interphase NSCs of MARCM clones in control (FRT82B) and *hyx^HT622^* were labelled for γ-tub, Asl, GFP, and PH3. γ-tub delocalization: control, 3.8%, n=26; *hyx^HT622^*, 95.7%, n=23; *hyx^w12-46^*, 54.8%, n=42. (B) Metaphase NSCs of MARCM clones in control (FRT82B) and *hyx^HT622^* were labelled for γ-tub, Asl, GFP, and PH3. γ-tub delocalization: control, 6.9%, n=29; *hyx^HT622^*, 93.1%, n=29; *hyx^w12-46^*, 70.8%, n=24. (C) Quantification graph of the fold change of γ-tub intensity in NSCs from A and B. (D) Interphase NSCs of MARCM clones in control (FRT82B) and *hyx^HT622^* were analyzed by Msps, Asl, GFP, and PH3. Control, n=44; *hyx^HT622^*, n=44; *hyx^w12-46^*, n=7. (E) Metaphase NSCs of control (FRT82B; all NSCs have Msps localization at the centrosomes, n=32), *hyx^HT622^* (n=24), and *hyx^w12-46^* (n=5) MARCM clones were labelled for Msps, Asl, GFP, and PH3. (F) Quantification graph of the fold change of Msps intensity in NSCs from D and E. Interphase: control, 1-fold, n=44; *hyx^HT622^*, 0.14-fold, n=44; *hyx^w12-46^*, 0.66-fold, n=7. Metaphase: control, 1-fold, n=32; *hyx^HT622^*, 0.19-fold, n=24; *hyx^w12-46^*, 0.48-fold, n=5. Cell outlines are indicated by white dotted lines. Arrows point at the centrosomes. Scale bars: 5 μm.

Next, we examined the localization of CNN, another essential PCM component. During interphase, 98.0% of control NSCs had intense CNN signal at the centrosomes marked by Asl (Figure S4D, F). By contrast, CNN was barely detectable at the centrosomes in 84.8% of *hyx^HT622^* and 58.3% of *hyx^w12-46^* (n=12) interphase NSCs (Figure S4D, F). Consistent with this, CNN levels were significantly diminished at the centrosomes in 72.4% of *hyx^HT622^* and 66.7% of *hyx^w12-46^* (n=18) metaphase NSCs (Figure S4E-F). Similarly, CNN levels were dramatically reduced from the centrosomes in 92.6% of interphase NSCs upon *hyx* knockdown (Figure S4G, I). In metaphase NSCs with *hyx* knockdown, CNN intensity at the centrosomes significantly dropped in 46.7% of NSCs (Figure S5H-I). Our observations indicate that Hyx ensures the recruitment of PCM proteins γ-tub and CNN to the centrosomes in both interphase and mitotic NSCs.

### Centrosomal localization of Msps, D-TACC, and Polo is dependent on Hyx function in NSCs

Mini spindles (Msps) is an XMAP215/ch-TOG family protein and a key microtubule polymerase that controls microtubule growth and asymmetric division of NSCs in *Drosophila* larval central brains (Lee, Gergely et al. 2001, Chen, Koe et al. 2016). In control interphase NSCs, Msps co-localized with Asl at the centrosomes (Figure 5D). However, during interphase, Msps was delocalized from the centrosomes in 86.4% of *hyx^HT622^* and 42.9% of *hyx^w12-46^* NSCs (Figure 5D). Likewise, during metaphase, Msps was nearly absent at the centrosomes in 87.5% of *hyx^HT622^* and 60.0% of *hyx^w12-46^* NSCs (Figure 5E). Moreover, Msps was undetectable at the centrosomes in the majority of interphase NSCs upon *hyx* knockdown (Figure S5A, C). In metaphase NSCs, Msps was detected at the centrosomes only in 16.4% of *hyx* RNAi NSCs (Figure S5B-C).

As the efficient centrosomal localization of Msps depends on D-TACC, a microtubule-binding centrosomal protein (Lee, Gergely et al. 2001), we wondered whether D-TACC localization in NSCs requires Hyx function. In all control interphase NSCs, D-TACC was concentrated at the centrosomes marked by Asl (Figure S5D). Remarkably, during interphase, D-TACC was absent from the centrosomes in 92.0% of *hyx^HT622^* (Figure S5D) and 81.8% of *hyx^W12-46^* NSCs (n=22). Similarly, in metaphase NSCs, D-TACC was undetectable at the centrosomes in 95.2% of *hyx^HT622^* NSCs (Figure S5E) and 80.0% of *hyx^w12-46^* NSCs (n=15). Likewise, D-TACC was apparently undetectable at the centrosomes in the majority of interphase and metaphase NSCs upon *hyx* knockdown (Figure S5G-H). The fluorescence intensity of D-TACC was significantly decreased in *hyx*-depleted NSCs (Figure S5F-I). Taken together, our results suggest that Hyx is essential for the centrosomal localization of Msps and D-TACC in cycling NSCs.

As Polo, another key centrosomal protein, is critical for the assembly of interphase microtubule asters and asymmetric cell division (Rusan and Peifer 2007, Wang, Ouyang et al. 2007), we tested whether Hyx regulates the localization of Polo at the centrosomes. In control interphase NSCs, Polo was strongly detected at the centrosome marked by Asl (Figure 6A). In contrast, Polo was almost completely absent in 86.8% of *hyx^HT622^* and 58.8% of *hyx^w12-46^* interphase NSCs (Figure 6A). Furthermore, in control metaphase NSCs, Polo mainly appeared on the centrosomes and kinetochores, and weakly on the mitotic spindle (Figure 6B). However, 80.0% of *hyx^HT622^* and 75% of *hyx^w12-46^* metaphase NSCs lost Polo loci, and the remaining NSCs only had a weak Polo signal (Figure 6B). Similarly, upon *hyx* knockdown, Polo was almost completely lost from the centrosomes in 84.6% of interphase NSCs and 78.3% of metaphase NSCs (Figure S6A-B). The fluorescence intensity of Polo was significantly reduced at the centrosomes in *hyx*-depleted interphase and metaphase NSCs (Figure 6C, S6C).

**Figure 6.**
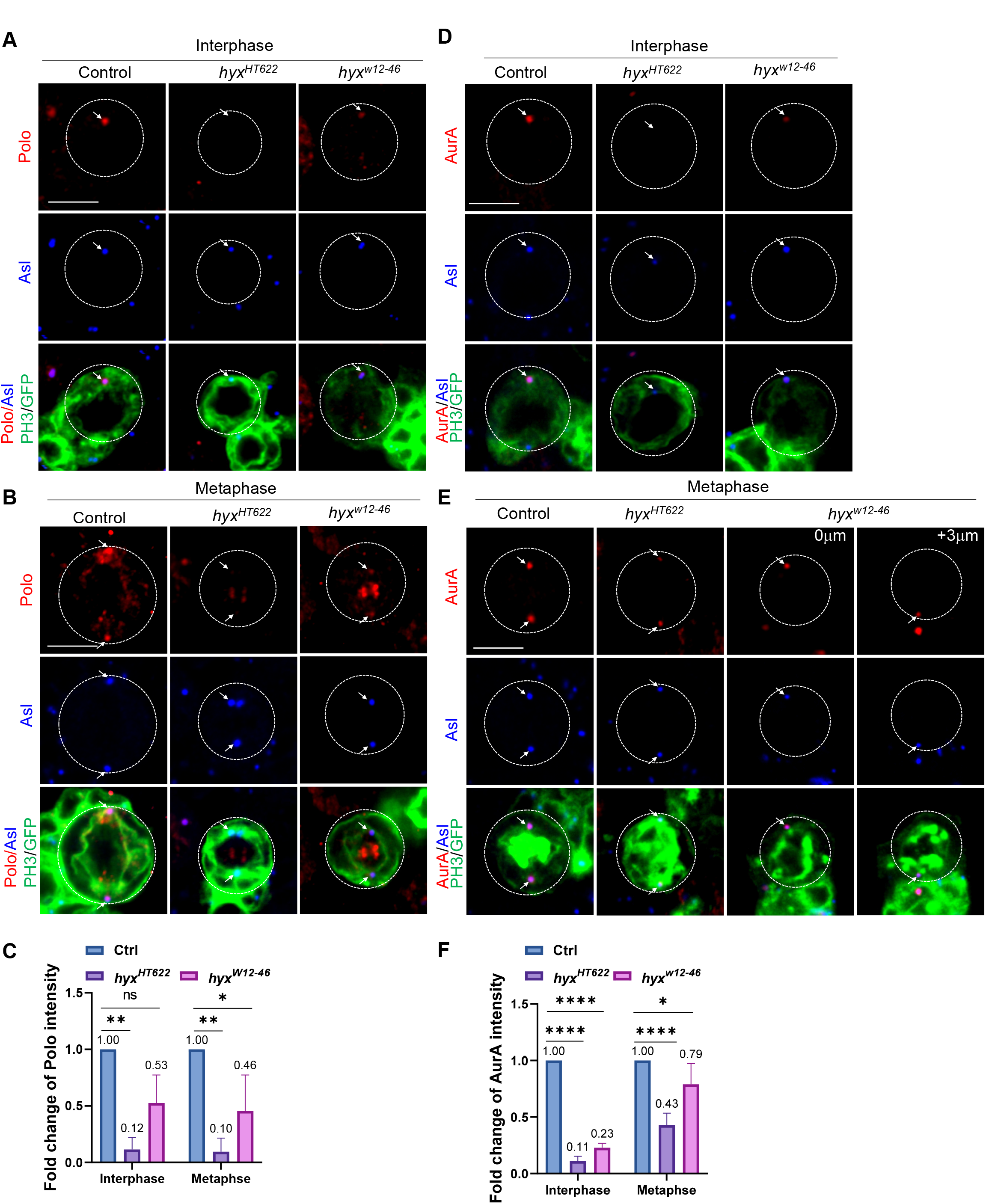
Hyx regulates the localization of centrosomal proteins Polo and AurA in NSCs. (A) Interphase NSCs from control (FRT82B) and *hyx^HT622^* MARCM clones were labelled for Polo, Asl, GFP, and PH3. Polo delocalization/reduction: control, 5.8%, n=52; *hyx^HT622^*, 86.8%, n=38; *hyx^w12-46^*, 58.8%, n=34. (B) Metaphase NSCs of control (FRT82B), *hyx^HT622^* and *hyx^w12-46^* MARCM clones were labelled for Polo, Asl, GFP and PH3. Polo delocalization/reduction: control, 0%, n=20; *hyx^HT622^*, 80.0%, n=10; *hyx^w12- 46^*, 75.0%, n=12. (C) Quantification graph of the fold change of Polo intensity in NSCs from A and B. Interphase: control, 1-fold, n=41; *hyx^HT622^*, 0.12-fold, n=38; *hyx^w12-46^*, 0.53-fold, n=34. Metaphase: control, 1-fold, n=20; *hyx^HT622^*, 0.10-fold, n=10; *hyx^w12-46^*, 0.46-fold, n=12. (D) Interphase NSCs from control (FRT82B; n=36) and *hyx^HT622^* (n=39) MARCM clones were labelled for AurA, Asl, GFP, and PH3. (E) Metaphase NSCs of control (FRT82B; n=22) and *hyx^HT622^* (n=18) MARCM clones were labelled for AurA, Asl, GFP, and PH3. (F) Quantification graph of the fold change of AurA intensity in NSCs from D and E. Interphase: control, 1-fold, n=36; *hyx^HT622^*, 0.11-fold, n=39; *hyx^w12- 46^*, 0.23-fold, n=20. Metaphase: control, 1-fold, n=22; *hyx^HT622^*, 0.43-fold, n=18; *hyx^w12- 46^*, 0.79-fold, n=8. Cell outlines are indicated by white-dotted lines. Arrows point at the centrosome in A. Scale bars: 5 μm.

Centrosomal protein AurA inhibits NSC overgrowth and regulates centrosome functions by directing the centrosomal localization of D-TACC and Msps (Giet, McLean et al. 2002, Wang, Somers et al. 2006). We sought to examine whether the centrosomal localization of AurA is dependent on Hyx. AurA is clearly observed at the centrosomes marked by Asl in control interphase and metaphase NSCs (Figure 6D-E). Remarkably, AurA was nearly undetectable at the centrosomes in 87.2% of *hyx^HT622^* and 75.0% *hyx^w12-46^* interphase NSCs (Figure 6D). The fluorescence intensity of AurA decreased to 0.11-fold in *hyx^HT622^* and 0.23-fold in *hyx^w12-46^* interphase NSCs (Figure 6D, F). Likewise, AurA levels were significantly reduced in 88.9% of *hyx^HT622^* and 50% of *hyx^w12-46^* metaphase NSCs (Figure 6E-F). Furthermore, AurA was diminished in 100% of interphase NSCs and 56.7% of metaphase NSCs upon *hyx* knockdown (Figure S6D-F). Taken together, our data show that Hyx plays an essential role in centrosome assembly and functions by recruiting major centrosomal proteins to the centrosomes in NSCs.

### Hyx is required for centrosome assembly in S2 cells *in vitro*

To investigate whether Hyx plays a role in centrosome assembly in non-neuronal cells, we knocked down *hyx* in S2 cells by dsRNA treatment. We found that a centriolar protein, Ana2, remained localized at the centrosomes in metaphase cells (Figure S6G). This suggests that Hyx is not essential for the localization of centriolar proteins in both S2 cells and NSCs. Next, we examined the localization of other centrosomal proteins in S2 cells. Remarkably, D-TACC intensity was significantly decreased at the centrosomes, upon *hyx* knockdown, in metaphase S2 cells (Figure 7A-B). Consistent with these observations, the intensity of α-tub was also decreased by 0.65-fold on mitotic spindles (Figure 7C-D). These *in vitro* data support our observations in the larval brain and indicate that Hyx regulates microtubule growth and the localization of centrosomal proteins. Polo is undetectable in interphase S2 cells, unlike its robust localization in NSCs during the interphase. Consistent to our *in vivo* observations, we found that the overall intensity of Polo was significantly reduced to 0.67-fold in the dividing metaphase cells upon *hyx* knockdown (Figure 7A, E). Also, γ-tub intensity at the centrosomes marked by Ana2 was similar to that observed in the control (Figure S6G-H). The different observations in S2 cells and larval brains are likely due to incomplete depletion of *hyx* in S2 cells and/or different underlying mechanisms *in vitro*.

**Figure 7.**
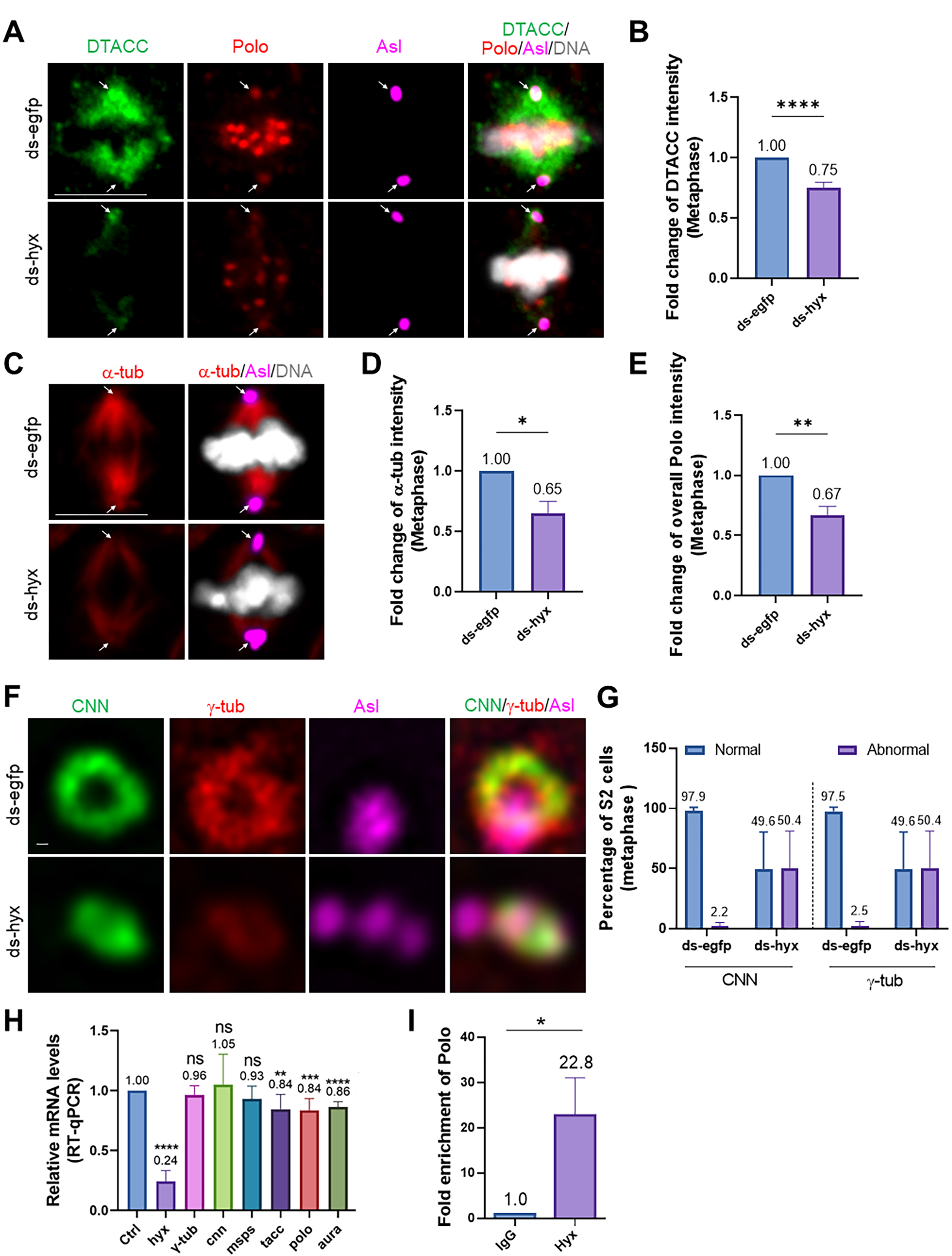
Hyx is required for centrosome assembly in S2 cells *in vitro*. (A) Metaphase cells from ds-*egfp*-treated S2 cells and ds-*hyx*-treated S2 cells were labeled for DTACC, Polo, Asl, and DNA. (B) Quantification graph of the fold change of DTACC intensity in S2 cells from A. ds-*egfp*, 1-fold, n=66; ds-*hyx*, 0.75-fold, n=69. (C) Metaphase cells from S2 cells treated with ds-*egfp* (n=83) and S2 cells treated with ds-*hyx* (n=88) were labeled for α-tub, Asl, and DNA. (D) Quantification graph of the fold change of α-tub intensity in S2 cells from C. (E) Quantification graph of the fold change of overall Polo intensity in S2 cells from A. ds-*egfp*, 1-fold, n=77; ds-*hyx*, 0.67-fold, n=81. (F) Spinning disc super-resolution imaging of CNN, γ-tub, and Asl in metaphase S2 cells treated with ds-*egfp* (CNN, n=46; γ-tub, n=37) and ds-*hyx* (CNN, n=39; γ-tub, n=39). (G) Quantification graph of the percentage of metaphase S2 cells forming “doughnut-like” shape of CNN and γ-tub in F. (H) Quantification graph of RT-qPCR for various genes in S2 cells from the control (ds-*egfp)* and ds-*hyx*. (I) Quantification graph of ChIP-qPCR for detecting *polo* occupancy by Hyx from IgG-immunoprecipitated DNA samples and Hyx-immunoprecipitated DNA samples in S2 cells. Arrows indicate the centrosomes. Scale bars: 5 μm.

To further probe how Hyx regulates centrosome assembly, we examined the ultrastructure of CNN and γ-tub using super-resolution imaging. CNN and γ-tub formed “doughnut-like” rings surrounding the centriolar protein Asl, at the centrosomes, in 97.9% and 97.5% of control metaphase cells, respectively (Figure 7F-G). Remarkably, CNN and γ-tub failed to form the ring patterns at the centrosomes in 50.4% of *hyx* knockdown mitotic cells; rather, they formed irregular shape at the centrosomes (Figure 7F-G). These observations suggest that Hyx is required for the proper recruitment of CNN and γ-tub at the centrosomes in S2 cells.

### Hyx directly regulates the expression of *polo*

As Parafibromin/Hyx regulates transcriptional events (Yang, Han et al. 2010), we wondered whether Hyx was required for the expression of genes that are involved in centrosome assembly. To this end, we performed reverse transcription quantitative real-time PCR (RT-qPCR) to detect differential transcription levels of those genes in S2 cells. Upon treatment of dsRNA against *hyx*, the relative mRNA level of *hyx* was dramatically decreased to 0.24-fold in comparison with the control 1-fold, suggesting that the knockdown was effective (Fig. 7H). The transcriptional levels of major centrosomal proteins, including DTACC, AurA, Polo, were significantly reduced in the S2 RNA extracts treated with dsRNA against *hyx* when comparing with control samples treated with dsRNA against *egfp* (Fig. 7H; *tacc*, 0.84-fold; *aura*, 0.86-fold; and *Polo*, 0.84-fold). The mRNA levels of *γ-tub*, *cnn*, and *msps* were not apparently affected upon *hyx* knockdown in S2 cells. Therefore, Hyx is required for the expression of centrosome-related genes.

To further investigate whether Hyx directly regulates the expression of *polo*, we performed Chromatin Immunoprecipitation (ChIP) coupled with quantitative PCR (ChIP-qPCR) and found a significant high enrichment of Hyx at potential *polo* promoter region compared with IgG control (Figure 7I, 22.8-fold *vs* 1.0-fold; 3 repeats), suggesting occupancy of *polo* by endogenous Hyx.

Taken together, Parafibromin/Hyx governs NSC polarity and centrosome assembly, and directly regulates the expression of *polo*.

## DISCUSSION

In this study, we established the essential role of Hyrax (Hyx), the *Drosophila* ortholog of Parafibromin, during the development of the central nervous system. We show that Hyx functions as a brain tumour suppressor-like protein by governing NSC asymmetric division and inhibiting NSC overgrowth in the central brains of *Drosophila* larvae. We also demonstrate that Hyx plays a novel function in the formation of microtubule asters and mitotic spindles in interphase NSCs. Particularly, Hyx is important for the localization of PCM proteins to the centrosomes in dividing NSCs and S2 cells. Therefore, ours is the first study to demonstrate that Parafibromin/Hyx has a tumour suppressor-like function and plays a critical role in the asymmetric division of NSCs, by regulating microtubule growth and centrosomal assembly in these cells.

It is established that *Drosophila* Hyx is essential for embryogenesis and wing development (Mosimann, Hausmann et al. 2006). In this study, we provide the first evidence that Hyx is crucial for *Drosophila* larval brain development. Furthermore, we showed that Hyx is essential for the polarized distribution of proteins in dividing NSCs, indicating a novel role for Hyx in regulating NSC apicobasal polarity. We also found that Hyx is required for the centrosomal localization of AurA and Polo kinases in NSCs, two brain tumor suppressor-like proteins that regulate asymmetric cell divisions (Lee, Andersen et al. 2006, Wang, Ouyang et al. 2007, Wang, Chang et al. 2009). Therefore, our study identifies a previously unknown link between Hyx and these cell cycle regulators, raising an interesting possibility that similar regulatory mechanisms may exist in other types of dividing cells, including cancer cells.

Human Parafibromin is a well-known tumour suppressor in parathyroid carcinomas and many other types of cancers (Selvarajan, Sii et al. 2008, Zheng, Takahashi et al. 2008, Newey, Bowl et al. 2009, Zhang, Yang et al. 2015). Parafibromin primarily regulates transcriptional events and histone modifications (Wei, Yu et al. 1999, Rea, Eisenhaber et al. 2000). It is also known to inhibit cell proliferation by the blockage of a G1 cyclin, Cyclin D1, and the c-myc proto-oncogene (Woodard, Lin et al. 2005, Lin, Zhang et al. 2008, Yang, Han et al. 2010). We show that Hyx controls *polo* expression likely through a direct transcriptional regulation.

In addition to its role in promoting asymmetric cell divisions and the establishment of apicobasal cell polarity, we provide compelling data that Hyx plays a novel role in regulating microtubule growth and centrosomal assembly in NSCs and S2 cells. Hyx was found to be important for the formation of interphase microtubule asters and the mitotic spindle. We also showed that it is required for the centrosomal localization of major PCM proteins, including γ-tub, CNN, AurA, and Polo. AurA is known to recruit γ-TuRC (γ-tubulin ring complex), CNN, and D-TACC to the centrosomes (Giet, McLean et al. 2002, Wang, Jiang et al. 2014). Therefore, Hyx might regulate the centrosomal recruitment of D-TACC, Msps, γ-tub, and CNN through AurA. Intriguingly, Hyx appeared to prevent centriole overduplication, suggesting it may play a pleiotropic function in regulating the cell cycle. The new role for Hyx in regulating microtubule growth and asymmetric divisions in NSCs, proposed in this study, supports our previous finding that microtubules play an essential role in NSC polarity (Chen, Koe et al. 2016). In addition to its predominant localization and functions in the nucleus, Parafibromin is also known to exist in the cytoplasm, where it regulates apoptosis by directly targeting p53 mRNA (Jo, Chung et al. 2014). Human Parafibromin directly interacts with actin-binding proteins, actinin-2 and actinin-3, during the differentiation of myoblasts (Agarwal, Simonds et al. 2008), suggesting that Parafibromin might regulate the actin cytoskeleton. Interestingly, *C. elegans* Ctr9 is required for the microtubule organization in epithelial cells during morphogenesis of the embryo (Kubota, Tsuyama et al. 2014). Therefore, the function of Hyx/Parafibromin in regulating microtubule growth and centrosomal assembly is likely a general paradigm in cell division regulation, which might be disrupted in cancer cells.

Parafibromin/HRPT-2 is expressed in both mouse and human brains (Porzionato, Macchi et al. 2006). Deletion of *Hrpt2* in mouse embryos results in early lethality and a developmental defect of the brain, suggesting that Parafibromin may play a role in CNS development (Wang, Bowl et al. 2008). Further investigations on the likely conserved functions of mammalian Parafibormin in NSC divisions and microtubule growth is warranted in future studies.

## ACKNOWLEDGMENTS

We thank Dr. Yu F. for providing the EMS mutant collection (on 3R chromosome) from which *hyx* alleles were isolated. We also thank Konard Basler, J. A. Knoblich, M. Buszczak, J. T. Lis, F. Matsuzaki, W. Chia, X. Yang, J. Skeath, F. Yu, YN Jan, C. Gonzalez, J. Raff, E. Schejter, T. Megraw and C. Sunkel, the Bloomington Drosophila Stock Center, the Vienna Drosophila Resource Center, the Kyoto stock centre DGGR, and the Developmental Studies Hybridoma Bank for fly stocks and antibodies. This work is supported by Singapore Ministry of Education Tier 2 MOE2018-T2-2-047 to H.W. and Spanish Ministerio de Ciencia, Innovacion y Universidades (PGC2018-097372-B-100) to C.G.

## AUTHOR CONTRIBUTIONS

Conceptualization, H.W., Q.D., and C.W.; Methodology, Data Curation, and Formal Analysis, Q.D., C.W., S.L., C.T., J.P.H., W-K.S., and Y.S.T.; Writing-Original draft, Q.D., C.W., and H.W.; Writing-Review & Editing, H.W., Q.D., and C.G.; Funding Acquisition, H.W. and C.G.; Resources, H.W.; Supervision, H.W.

## METHODS

### Fly stocks and genetics

Fly stocks and genetic crosses were reared at 25°C unless otherwise stated. Fly stocks were kept in vials or bottles containing standard fly food (0.8% *Drosophila* agar, 5.8% Cornmeal, 5.1% Dextrose, and 2.4% Brewer’s yeast). The following fly strains were used in this study: *insc-*Gal4 (BDSC#8751; 1407-Gal4), *insc-*Gal4, *UAS-Dicer2* with and without *UAS-CD8-GFP*, *Jupiter-GFP* (G147), UAS*-hyx*, *UAS-HRPT2* (Mosimann, Hausmann et al. 2006). The type II NSC driver: w; *UAS-Dicer 2*, *wor*-Gal4, *ase*-Gal80/CyO; *UAS-mCD8-GFP*/TM3, Ser (Neumüller, Richter et al. 2011).

The following stocks were obtained from Bloomington Drosophila Stock Center (BDSC): *UAS-Gal* RNAi (BDSC#50680; this stock is often used as a control UAS element to balance the total number of UAS elements), *ctr9^12P023^* (BDSC#59389), *rtf1* RNAi (BDSC#34586), *rtf1* RNAi (BDSC#34850), *rtf1* RNAi (BDSC#31718).

The following stocks were obtained from Vienna Drosophila Resource Center (VDRC): *hyx* RNAi (28318), *hyx* RNAi (103555), *atms* RNAi (108826), *atu* RNAi (17490), *atu* RNAi (106074), *ctr9* RNAi (108874), *ctr9* RNAi (Chaturvedi, Inaba et al. 2016), *rtf1* RNAi (27341), *rtf1* RNAi (110392).

All experiments were carried out at 25°C, except for RNAi knockdown or overexpression experiments that were performed at 29°C.

### Immunohistochemistry

Third-instar *Drosophila* larvae were dissected in PBS, and larval brains were fixed in 4% EM-grade formaldehyde in PBT (PBS + 0.3% Triton-100) for 22 min. The samples were processed for immunostaining as previously described (Li, Koe et al. 2017). For α-tubulin immunohistochemistry, larvae were dissected in Shield and Sang M3 medium (Sigma-Aldrich), supplemented with 10% FBS, followed by fixation in 10% formaldehyde in Testis buffer (183 mM KCl, 47 mM NaCl, 10 mM Tris-HCl, and 1 mM EDTA, pH 6.8), supplemented with 0.01% Triton X-100. The fixed brains were washed once in PBS and twice in 0.1% Triton X-100 in PBS. Images were taken with an AxioCam HR camera (with 1.5× to 8× of digital zoom) of a LSM710 confocal microscope system (Axio Observer Z1; ZEISS), using a Plan-Apochromat 40×/1.3 NA oil differential interference contrast objective. The brightness and contrast of the images obtained were adjusted using Adobe Photoshop or Fiji (imageJ).

The primary antibodies used were: rabbit affinity-purified anti-Hyx/Cdc73 (1: 1000; J. T. Lis); guinea pig anti-Dpn (1:1000), mouse anti-Mira (1:50; F. Matsuzaki), rabbit anti-Mira (1:500, W. Chia), anti-Insc (1:1000, X. Yang), rabbit anti-aPKCζ C20 (1:100; Santa Cruz Biotechnologies, Dallas, TX), guinea-pig anti-Numb (1:1000; J Skeath), rabbit anti-GFP (1:3,000; F. Yu), mouse anti-GFP (1:5,000; F. Yu), rabbit anti-Asense (1:1000; YN Jan), guinea pig anti-Asl (1:200, C. Gonzalez), rabbit anti-Sas-4 (1:100, J. Raff), mouse anti-α-tubulin (1:200, Sigma, Cat#: T6199), mouse anti-γ-tubulin (1:200, Sigma, Cat#: T5326), rabbit anti-CNN (1:5000, E. Schejter and T. Megraw), rabbit anti-Msps (1:500), rabbit anti-Msps (1:1000, J. Raff), rabbit anti-PH3 (1:200, Sigma, Cat#: 06-570), rabbit anti-DTACC (1:200), rabbit anti-Ana2 (Wang, Li et al. 2011), α-tubulin (1:200, Sigma, Cat#: T6199), rabbit anti-AurA (1:200, J. Raff), rat anti-CD8 (1:250, Caltag laboratories), mouse anti-Polo (1:30, C. Sunkel). The secondary antibodies used were conjugated with Alexa Fluor 488, 555, or 647 (Jackson laboratory).

### Spinning disc super-resolution imaging

Super-resolution Spinning Disc Confocal-Structured Illumination Microscopy (SDC-SIM) was performed on a spinning disk system (Gataca Systems) based on an inverted microscope (Nikon Ti2-E; Nikon) equipped with a confocal spinning head (CSU-W; Yokogawa), a Plan-Apo objective (100x1.45-NA), and a back-illuminated sCMOS camera (Prime95B; Teledyne Photometrics). A super-resolution module (Live-SR; GATACA Systems) based on structured illumination with optical reassignment technique and online processing leading to a two-time resolution improvement (Roth and Heintzmann 2016) is included. The maximum resolution is 128 nm with a pixel size of 64nm in super-resolution mode. Excitation light at 488-nm/150mW (Vortran) (for GFP), 561-nm/100mW (Coherent) (for mCherry/mRFP/tagRFP) and 639-nm/150mW (Vortran) (for iRFP) was provided by a laser combiner (iLAS system; GATACA Systems), and all image acquisition and processing were controlled by the MetaMorph (Molecular Device) software. Images were further processed with imageJ.

### Generation of guinea pig anti-Hyx antibodies

The cDNA region encoding the N-terminal 1-176 amino acid residues of Hyx/Cdc73 was amplified by PCR with the oligos: 5’-TCCGAATTCATGGCAGATCCGCTCA GC-3’and 5’-ATGCGGCCGCCTACGTCTCGGACAGCGACTT-3’. The PCR products were cloned into the pMAL-c2x (Addgene # 75286) vector. The fusion protein MBP-Hyx 1-176 was purified and injected into guinea pigs and the antibodies generated were purified by GenScript (Hong Kong).

### Clonal analysis

MARCM clones were generated as previously described (Lee and Luo 1999). Briefly, larvae were heat-shocked at 37°C for 90 min at 24 hr after larval hatching (ALH) and 10–16 hr after the first heat shock. Larvae were further aged for 3 days at 25°C, and larval brains were dissected and processed for immunohistochemistry. To generate type II NSC clones, UAS lines were crossed to the type II driver at 25°C and shifted to 29°C at 24 hr ALH. Wandering third instar larvae were dissected after incubation for 3 or 4 days at 29°C.

### Time-lapse recording

Time-lapse recording was performed as described (Chen, Koe et al. 2016). The whole-mount brain expressing G147-GFP was used to analyze the asymmetric cell division of NSCs. The brain was dissected and loaded into a Lab-TekTM chambered coverglass, (Thermo Fisher Scientific) filled with dissecting medium that is supplemented with 2.5% methyl cellulose (Sigma-Aldrich). The time-lapse images of NSC divisions were acquired every 30 s on a confocal microscope (LSM 710; ZEISS). The video was processed with ImageJ and displayed at 15 frames per second.

### Microtubule regrowth assay

Microtubule regrowth assay was performed as described previously (Chen, Koe et al. 2016). Third-instar larval brains were dissected in Shield and Sang M3 insect medium (Sigma-Aldrich) supplemented with 10% FBS, and microtubules were depolymerized by incubating the larval brains on ice for 40 min. The brains were allowed to recover at 25°C for various time periods to facilitate microtubule regrowth. The brains were immediately fixed in 10% formaldehyde in testis buffer (183 mM KCl, 47 mM NaCl, 10 mM Tris-HCl, and 1 mM EDTA, pH 6.8) supplemented with 0.01% Triton X-100. The fixed brains were washed once in PBS and twice in 0.1% Triton X-100 in PBS, following which they were processed for immunohistochemistry.

### S2 cell culture, transfection, and quantitative RT-PCR

#### Cell culture

*Drosophila* S2 cells were cultured in Express Five SFM (Thermo Fisher Scientific), supplemented with 2 mM glutamine (Thermo Fisher Scientific), at 25°C.

#### dsRNA production

DNA fragments, approximately 470 bp in length for dsEGFP as control and 825bp in length for dsHyx, were amplified using PCR. Each primer used in the PCR contained a 5′ T7 RNA polymerase binding site (TAATACGACTCACTATAGGG) followed by sequences specific for the targeted genes. The PCR products were purified by using the QIAquick PCR Purification Kit (Cat No. 28106). The purified PCR products were used as templates for the synthesis of dsRNA, by using a MEGASCRIPT T7 transcription kit (Ambion, Austin, TX). The dsRNA products were ethanol-precipitated and resuspended in water. The dsRNAs were annealed by incubation at 65°C for 30 min followed by slow cooling to room temperature. One microgram of dsRNA was analyzed by 1% agarose gel electrophoresis to ensure that the majority of the dsRNA existed as a single band. The dsRNA was stored at −20°C.

The primers used for dsRNA synthesis were:

ds-*egfp*-forward: 5’-TCGTGACCACCCTGACCTAC-3’;

s-*egfp*-reverse: 5’-GCTTCTCGTTGGGGTCTTT-3’;

ds-*hyx*-forward: 5’-TGCTGCAACACTCGGTCTAC-3’;

ds-*hyx*-reverse: 5’-GTGCTCCCGGTAGGTTGTTA- 3’.

#### Extraction of total messenger RNA (mRNA) and RT-qPCR

Total mRNA from ds- *egfp* and ds-*hyx*-treated S2 cells were extracted using TRI Reagent (Sigma-Aldrich) according to the manufacturer’s instructions. Reverse transcription was performed with iScript™ cDNA Synthesis Kit (Bio-RAD) according to the manufacturer’s instructions. RT-qPCR was performed according to the manufacturer’s instructions (SsoFast™ EvaGreen®, Bio-RAD).

The primers pairs used for RT-qPCR were:

*hyx* forward: 5’-AGCCGGCTCGAATAGCCAAAC-3’;

*hyx* reverse: 5’- TGAGCATGGTAATGAGGCTTG-3’;

*γ-tub23C* forward: 5’- ACCGCAAGGATGTGTTCTTC-3’;

*γ-tub23C* reverse: 5’- CCTCCGTGCTTGGATAGGTA-3’;

*cnn* forward: 5’-CCGGCAGGATATCTAGCGTA-3’;

*cnn* reverse: 5’-TTGCTGTCCGGTGATGTAGA-3’;

*msps* forward: 5’-TTACGCGACCAAATGATGAC-3’;

*msps* reverse: 5’-TACACACCAGCGCCTTACTG-3’;

*tacc* forward: 5’-AGCACTTGCAAGCCATGAGT-3’;

*tacc* reverse: 5’-GCCTTCTGTTGATCCATGCT-3’;

*polo* forward: 5’-AGAGCCTGTACCAGCAGCTC-3’;

*tacc* reverse: 5’-CTGCAGGATCTGTGTTCTCG-3’;

*aurA* forward: 5’- AAGAAGACCACATCAGAGTTTGC -3’;

*aurA* reverse: 5’- TTGATGTCCCTGTGTATGATGTC -3’.

### Chromatin immunoprecipitation

Chromatin immunoprecipitation (ChIP) was performed according to the manufacturer’s protocol (Millipore, 17-10085). Sonicated lysates were used for ChIP with antibodies against Hyx (J. T. Lis) and normal serum as a control. Immunoprecipitated DNA and input DNA were analyzed by quantitative real-time PCR using specific primers to polo promoter region: (5′-TACCAGAAAGTGTGCGATAGCC-3′ and 5′- GAAACGGAGATCAGATCCACAC -3′).

### Quantification and statistical analysis

*Drosophila* larval brains from various genotypes were placed dorsal side up on confocal slides. The confocal z-stacks were taken from the surface to the deep layers of the larval brains. For each genotype, at least 10 NSCs were imaged and ImageJ or Zen software was used for quantifications.

Statistical analysis was essentially performed using GraphPad Prism 8. Unpaired two-tail t-tests were used for comparison of two sample groups, and one-way ANOVA or two-way ANOVA followed by Sidak’s multiple comparisons test was used for comparison of more than two sample groups. All data are shown as the mean ± SD. Statistically nonsignificant (ns) denotes P > 0.05, * denotes P <0.05, ** denotes P <0.01, *** denotes P <0.001, and **** denotes P < 0.0001. All experiments were performed with a minimum of two repeats. In general, n refers to the number of NSCs counted unless otherwise indicated.

**Movie S1.** Time-lapse imaging of G147/+ (Jupiter-GFP) NSCs under the control of *insc*-Gal4 at 48h ALH larval brains. Time scale: minute: second. Scale bar: 5 µm.

**Movie S2.** Time-lapse imaging of *hyx* RNAi (KK/V103555); G147 NSCs under the control of *insc*-Gal4 at 48h ALH larval brains. Time scale: minute: second. Scale bar: 5 µm.

---

**Figure S1.**
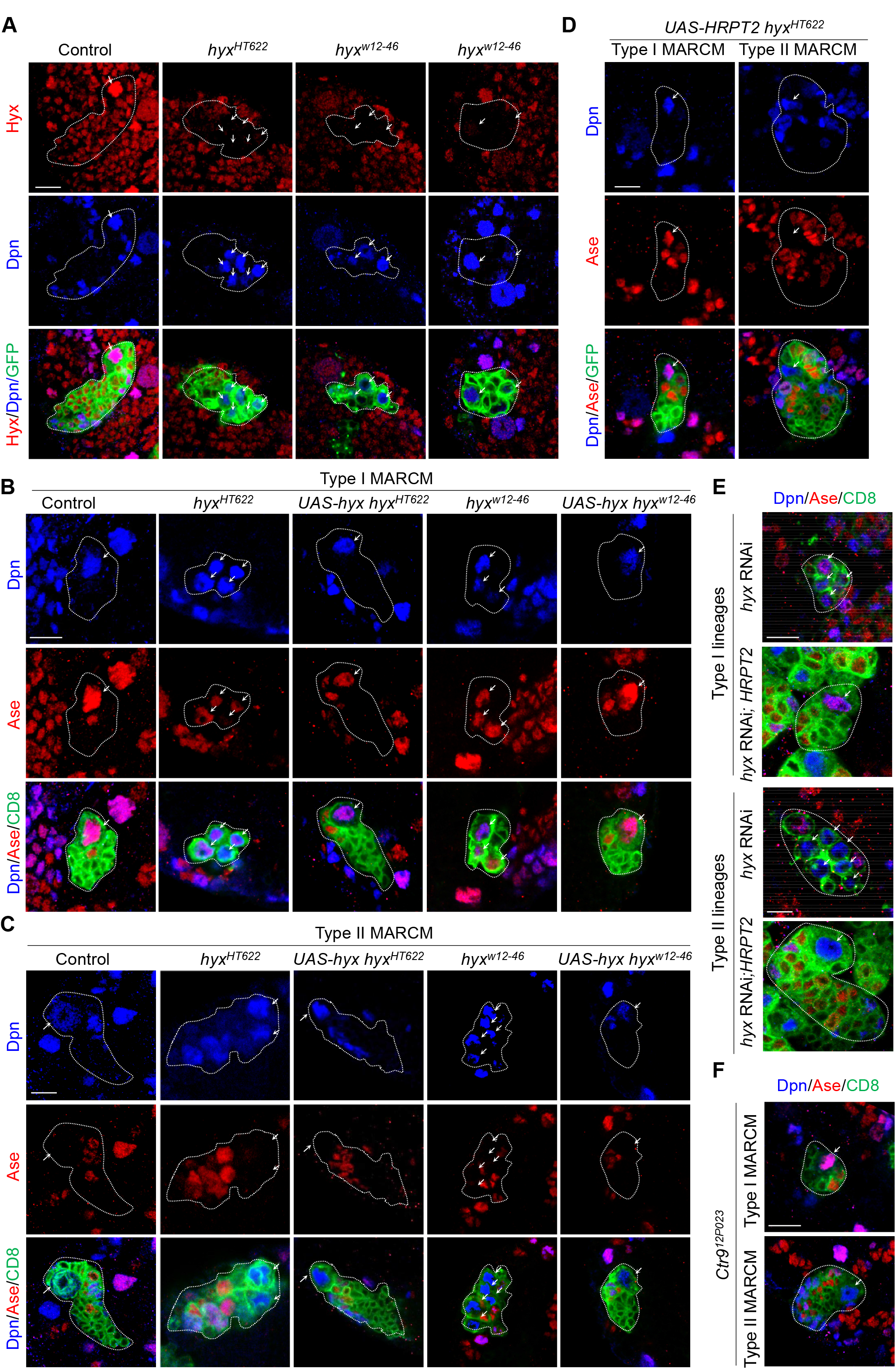
Hyx regulates NSC homeostasis of *Drosophila* larval central brains. (A) MARCM clones of control (FRT82B; n=20), *hyx^HT622^* (n=23), and *hyx^W12-46^* (n=25) were labelled for Hyx, Dpn, and GFP. (B) Type I MARCM clones of control (FRT82B; n=20), *hyx^HT622^* (n=17)*, hyx^W12-46^* (n=30), *UAS-hyx hyx^HT622^* (n=21), and *UAS-hyx hyx^W12- 46^* (n=40) were labelled for Dpn, Ase, and CD8. Ectopic NSCs were observed in 88.2% of *hyx^HT622^* and 36.7% of *hyx^w12-46^* larvae, but not in the control or rescued larvae. (C) Type II MARCM clones of control (FRT82B), *hyx^HT622^, hyx^W12-46^*, *UAS-hyx hyx^HT622^*, and *UAS-hyx hyx^W12-46^* were labelled for Dpn, Ase, and CD8. Ectopic NSCs were observed in *hyx^HT622^* (81.0%, n=21) and *hyx^w12-46^* (78.5%, n=30) larvae, but not in control (n=20), *UAS-hyx hyx^HT622^* (n=17) and *UAS-hyx hyx^W12-46^* (n=40) larvae. (D) MARCM clones of *UAS-HRPT2 hyx^HT622^* type I (n=30) and type II (n=7) were labelled for Dpn, Ase, and GFP. (E) Type I and type II NSC lineages from *hyx* RNAi (V103555 with *UAS-CD8-GFP*) and *UAS-HRPT2; hyx* RNAi under the control of *insc*-Gal4 driver were labelled for Dpn, Ase, and CD8 (n=10 brain lobes for each genotype). (F) Type I (n=12) and type II (n=16) MARCM clones from *ctr9^12P023^* were labelled for Dpn, Ase, and CD8. Clones are outlined with white dotted lines. NSCs and NSC-like cells are pointed by arrows. Scale bars: 5 μm.

**Figure S2.**
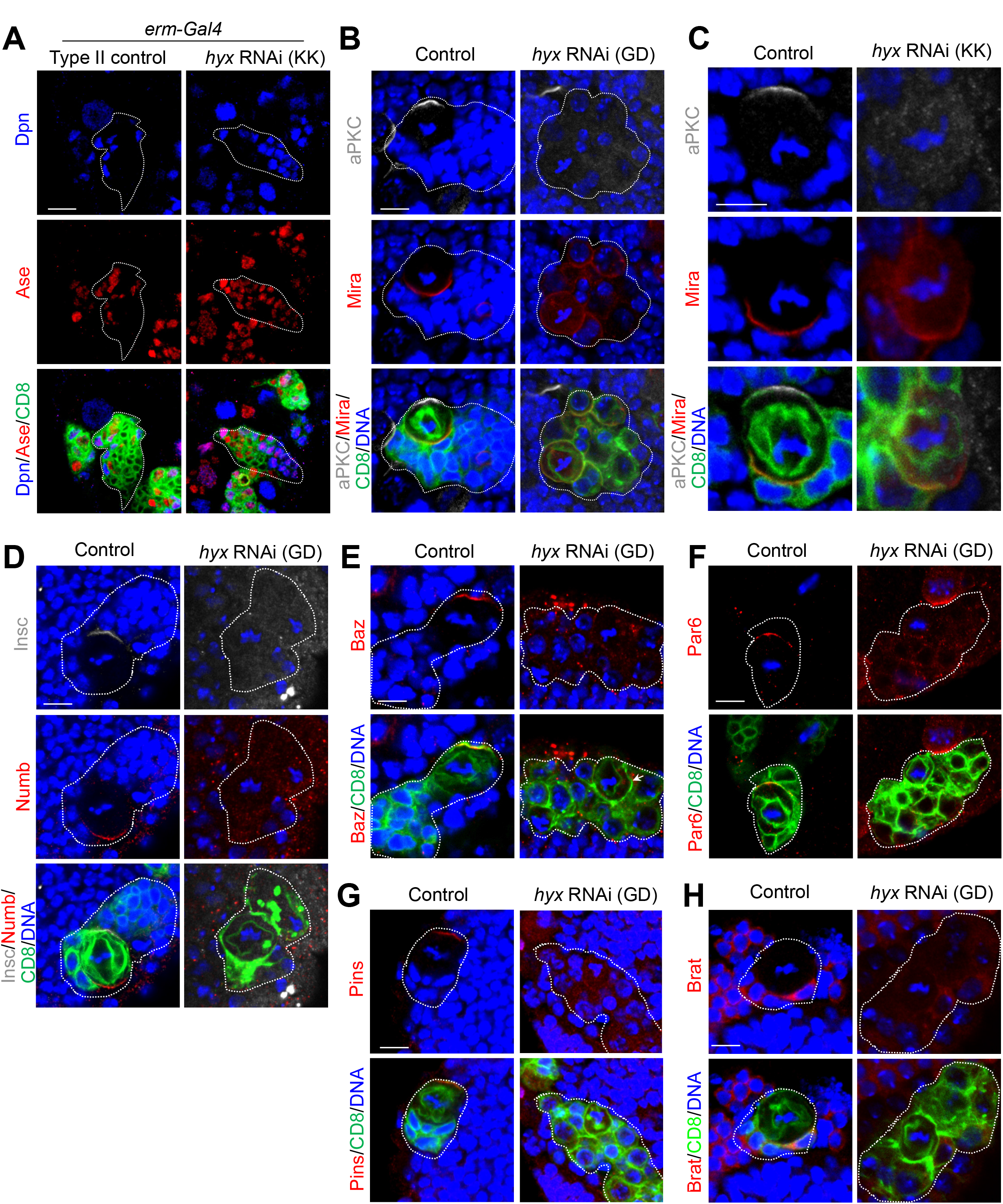
Hyx governs asymmetric cell division of NSCs. (A) INP lineages of control (*UAS-Dicer2*) and *hyx* RNAi (KK/V103555 with *UAS-Dicer2*) driven by *erm*-Gal4, *UAS-CD8-GFP* were labelled for Dpn, Ase, and CD8 (n=30 for both). (B) Metaphase NSCs of control (*insc*>*CD8-GFP*; n=50) and *hyx* RNAi (GD/V28318 with *UAS-CD8-GFP*) type II lineages were labeled for aPKC, Mira, CD8, and DNA. *hyx* RNAi: aPKC delocalization, 100%, n=50; Mira delocalization, 70%, n=50. (C) Metaphase NSCs from control (*insc>CD8-GFP*) and *hyx* RNAi (KK/V103555 with *UAS-CD8-GFP*) type II lineages were labeled for aPKC, Mira, CD8, and DNA. In *hyx* RNAi, delocalization of aPKC: 100%; Mira: 90%; n=50 for all. (D) Metaphase NSCs from control (*insc>CD8-GFP*) and *hyx* RNAi (GD/V28318 with *UAS-CD8-GFP*) type II lineages were labeled with Insc, Numb, CD8, and DNA. *hyx* RNAi, 100% delocalization of Insc and Numb; n=50 for all. (E) Metaphase NSCs from control (*insc>CD8-GFP*) and *hyx* RNAi (GD/V28318 with *UAS-CD8-GFP*) type II lineages were labeled for Baz, CD8, and DNA; *hyx* RNAi: 100% delocalization of Baz; n=50 for both genotypes. (F) Metaphase NSCs from control (*insc>CD8-GFP*) and *hyx* RNAi (GD/V28318 with *UAS-CD8-GFP*) type II lineages were labeled for Par6, CD8, and DNA. *hyx* RNAi: Par6 delocalization, 100%; n=50 for both genotypes. (G) Metaphase NSCs from control (*insc>CD8-GFP*) and *hyx* RNAi (GD/V28318 with *UAS-CD8-GFP*) type II lineages were labeled for Pins, CD8, and DNA. *hyx* RNAi: Pins delocalization, 100%; n=50 for both genotypes. (H) Metaphase NSCs from control (*insc>CD8-GFP*; n=50) and *hyx* RNAi (GD/V28318 with *UAS-CD8-GFP*) type II lineages were labeled for Brat, CD8, and DNA. *hyx* RNAi: Brat delocalization, 100%; n=50 for both genotypes. Clones are outlined by white-dotted lines. Scale bars: 5 μm.

**Figure S3.**
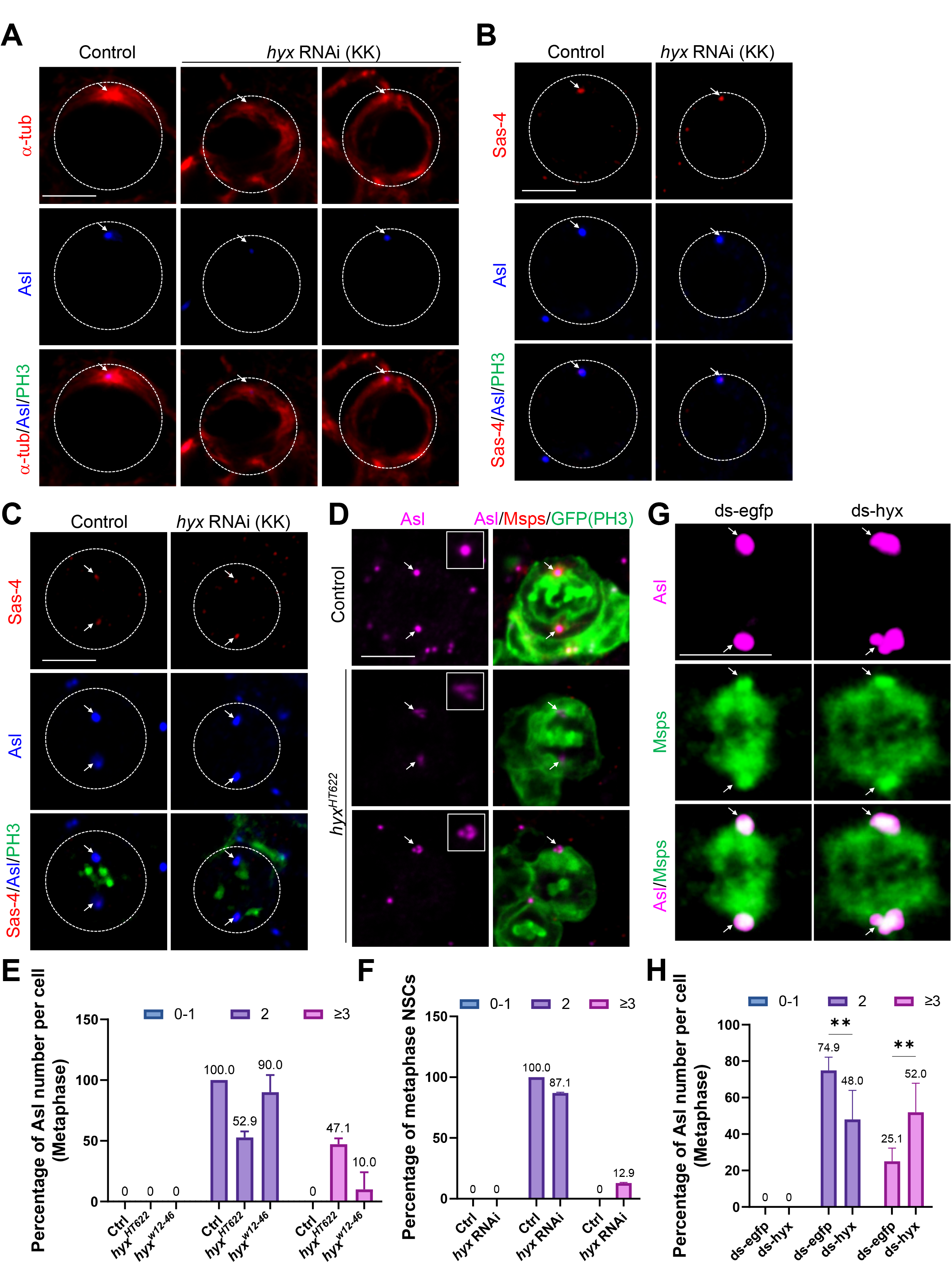
Hyx is required for the formation of microtubule aster and centriole number in NSCs. (A) Interphase NSCs from control (*UAS-β-Gal* RNAi; 100% aster formation, n=20) and *hyx* RNAi (KK/V103555; n=22) were labelled for α-tub, Asl, and PH3. (B) Interphase NSCs of control (*UAS-β-Gal* RNAi; n=23) and *hyx* RNAi (KK/V103555; n=20) were labelled for Sas-4, Asl, and PH3. (C) Prometa/metaphase NSCs of control (*UAS-β-Gal* RNAi) and *hyx* RNAi (KK/V103555) were labeled for Sas-4, Asl, and PH3 (n=23 for both). (D) Metaphase NSCs of control (FRT82B; n=54) and *hyx^HT622^* (n=37) were labelled for Asl, Msps, GFP, and PH3. Zoom-in areas are boxed. (E) Quantification graph showing the percentage of metaphase NSCs with indicated number of Asl per NSC in (D) and *hyx^w12-46^* (n=8). (F) Quantification graph showing the percentage of metaphase NSCs from control (*UAS-β-Gal* RNAi; n=31) and *hyx* RNAi (KK/V103555; n=23) with indicated number of Asl. (G) Metaphase ds-*egfp*-treated S2 cells (n=195) and ds-*hyx*-treated S2 cells (n=172) were labeled for Msps and Asl. (H) Quantification graph displaying the percentage of metaphase S2 cells with the indicated number of Asl per NSC in G. *hyx* knockdown was driven by *insc*-Gal4 in A-C. NSCs are circled by dotted lines. Arrows indicate the centrosomes. Scale bars: 5 μm.

**Figure S4.**
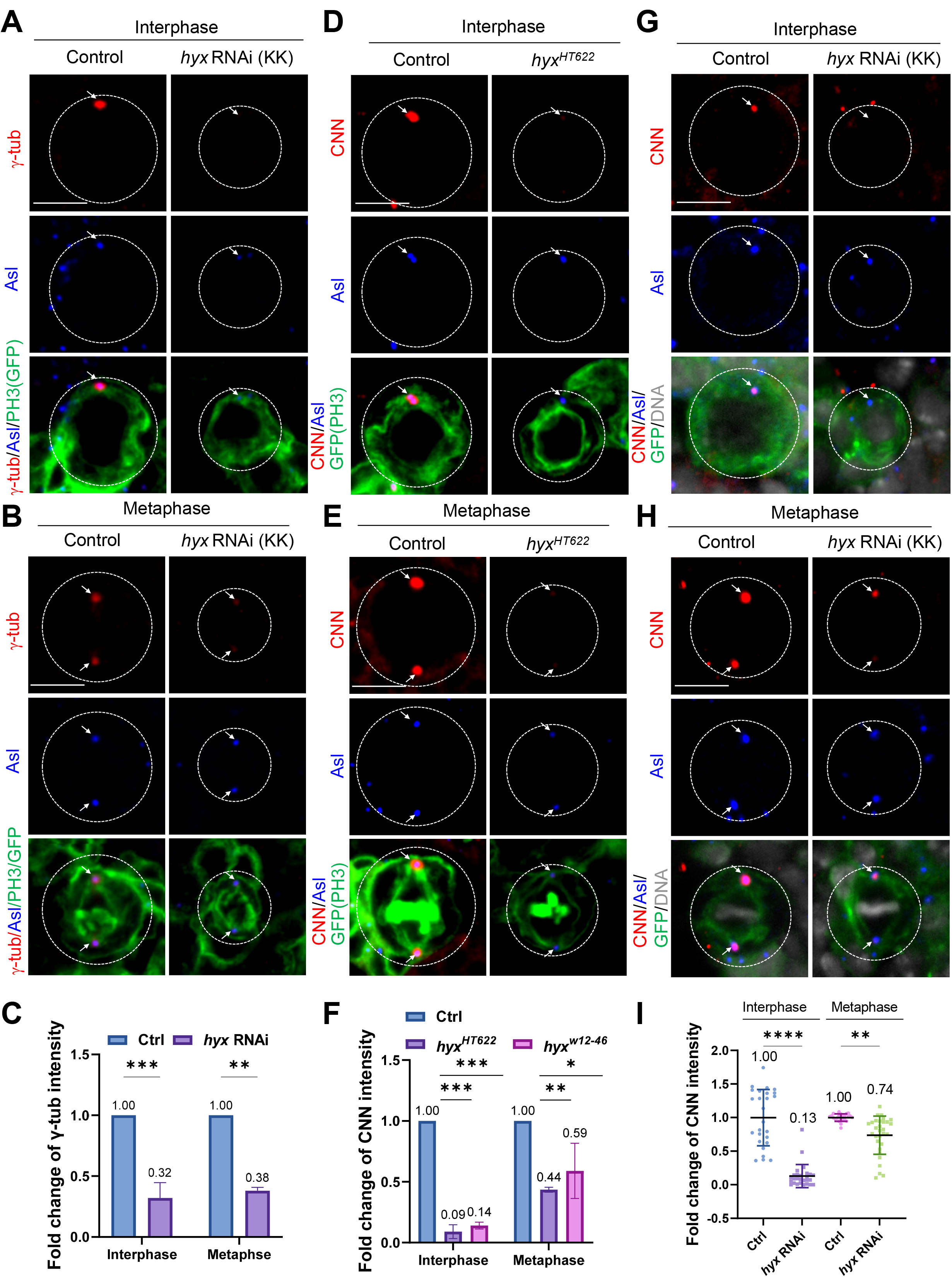
Hyx is required for the recruitment of PCM proteins γ-tub and CNN to the centrosome in NSCs. (A) Control (*UAS-β-Gal* RNAi; n=36) and *hyx* RNAi (KK/V103555; n=42) interphase NSCs were labelled for γ-tub, Asl, GFP, and PH3. (B) Control (*UAS-β-Gal* RNAi; n=25) and *hyx* RNAi (KK/V103555; n=32) metaphase NSCs were labelled for γ-tub, Asl, GFP, and PH3. In control, robust distribution of γ-tub was seen in 88.9% of interphase (A) and 92.0% of metaphase (B) NSCs. (C) Quantification graph of the fold change of γ-tub intensity in NSCs from A and B. Interphase: control, 1-fold, n=36; *hyx* RNAi, 0.32-fold, n=42. Metaphase: control, 1-fold, n=25; *hyx* RNAi, 0.38-fold, n=32. (D) Interphase NSCs of control (FRT82B; n=51) and *hyx^HT622^* (n=33) MARCM clones were labelled for CNN, Asl, GFP, and PH3. (E) Metaphase MARCM clones of control (FRT82B; all NSCs have robust CNN localization, n=27) and *hyx^HT622^* (n=29), were labelled for CNN, Asl, GFP, and PH3. (F) Quantification graph of the fold change of CNN intensity in NSCs from D and E. Interphase: control, 1-fold, n=51; *hyx^HT622^*, 0.09-fold, n=33; *hyx^w12-46^*, 0.14-fold, n=12. Metaphase: control, 1-fold, n=27; *hyx^HT622^*, 0.44-fold, n=29; *hyx^w12-46^*, 0.59-fold, n=18. (G) Interphase NSCs from control (*UAS-β-Gal* RNAi; 96.3% robust CNN signal) and *hyx* RNAi (KK/V103555) were labelled for CNN, Asl, GFP, and DNA (n=27 for both). (H) Metaphase NSCs of control (*UAS-β-Gal* RNAi; 100% CNN robust localization, n=24) and *hyx* RNAi (KK/V103555; n=30) were labelled for CNN, Asl, GFP, and DNA. (I) Quantification graph of the fold change of CNN intensity in NSCs from G and H. Interphase: control, 1-fold, n=27; *hyx* RNAi, 0.13-fold, n=27. Metaphase: control, 1-fold, n=24; *hyx* RNAi, 0.74-fold, n=30. *hyx* knockdown was driven by *insc*-Gal4 in A-I. NSCs are circled by dotted lines. Arrows indicate the centrosomes. Scale bars: 5 μm.

**Figure S5.**
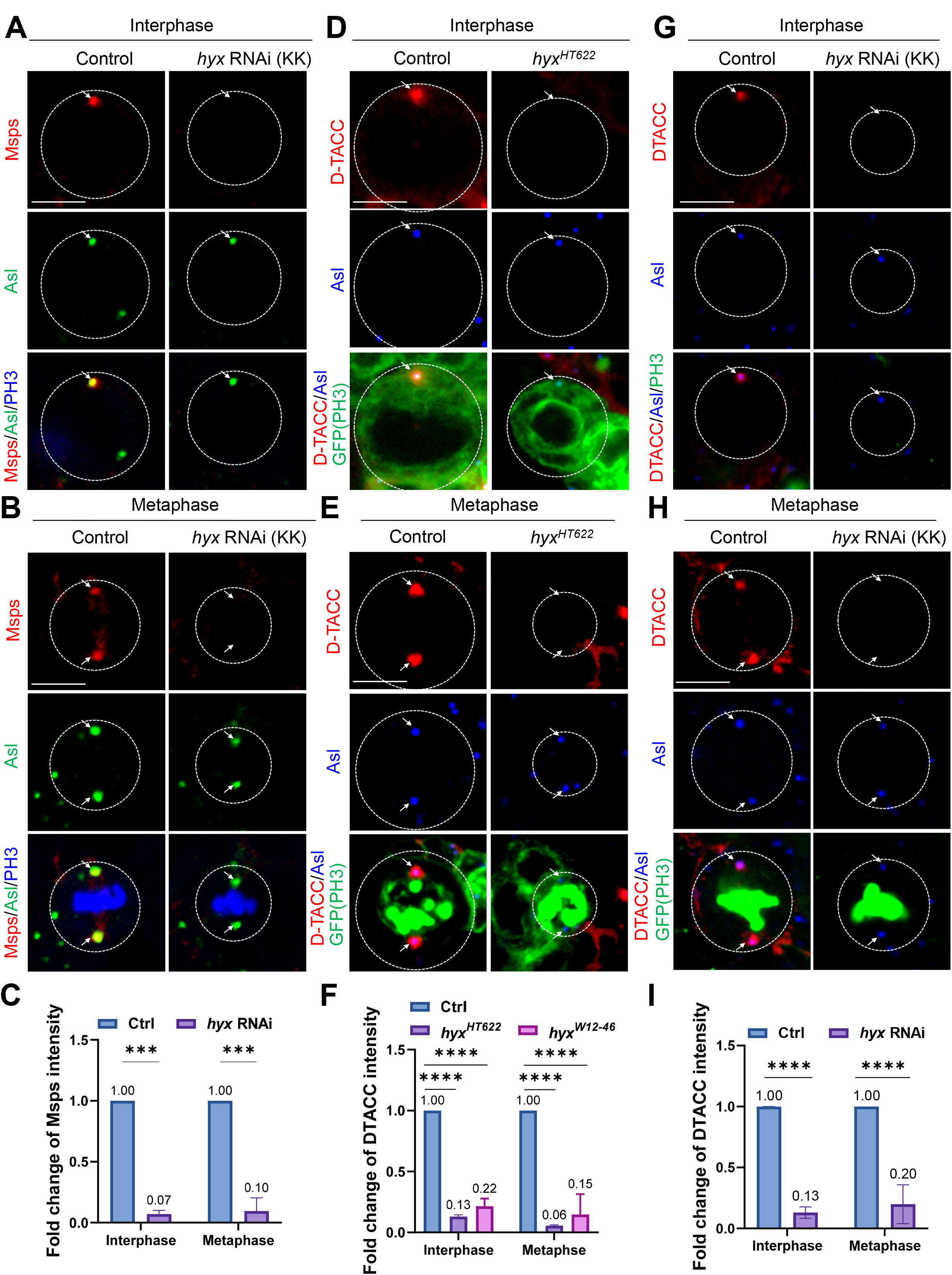
Msps and D-TACC were delocalized from the centrosomes upon *hyx* depletion in NSCs. (A) Interphase NSCs of control (*UAS-β-Gal* RNAi) and *hyx* RNAi (KK/V103555) were labelled for Msps, Asl, and PH3. Msps delocalization at the centrosomes: control: 0%, n=55; *hyx* RNAi, 82.5%, n=57. (B) Metaphase NSCs of control (*UAS-β-Gal* RNAi) and *hyx* RNAi (KK/V103555) were labelled for Msps, Asl, and PH3. Msps localization? at the centrosomes: control: 98.5%, n=55; *hyx* RNAi, 16.4%, n=67. (C) Quantification graph of the fold change of Msps intensity in NSCs from A and B. Interphase: control, 1-fold, n=25; *hyx* RNAi, 0.07-fold, n=25. Metaphase: control, 1-fold, n=25; *hyx* RNAi, 0.10-fold, n=26. (D) Interphase NSCs from control (FRT82B; n=41) and *hyx^HT622^* (n=25) MARCM clones were labelled for DTACC, Asl, GFP, and PH3. (E) MARCM clones of control (FRT82B; 100% D-TACC localization, n=18) and *hyx^HT622^* (n=21) were labelled for DTACC, Asl, GFP, and PH3. (F) Quantification graph of the fold change of DTACC intensity in NSCs from D and E. Interphase: control, 1-fold, n=21; *hyx^HT622^*, 0.13-fold, n=25; *hyx^w12-46^*, 0.22-fold, n=22. Metaphase: control, “1” fold, n=18; *hyx^HT622^*, “0.06”, n=21; *hyx^w12-46^*, “0.15” fold, n=15). (G) Interphase NSCs of control (*UAS-β-Gal* RNAi) and *hyx* RNAi (KK/V103555) were labelled for DTACC, Asl, and PH3. DTACC delocalization at the centrosomes: control, 3.8%, n=26; *hyx* RNAi, 83.3%, n=24. (H) Metaphase NSCs of control (*UAS-β-Gal* RNAi) and *hyx* RNAi (KK/V103555) were labelled for DTACC, Asl, and PH3. DTACC delocalization at the centrosomes: control, 3.3%, n=30; *hyx* RNAi, 85.3%, n=34. (I) Quantification graph of the fold change of DTACC intensity in NSCs from G and H. Interphase: control, 1-fold, n=26; *hyx* RNAi, 0.13-fold, n=24. Metaphase: control, 1-fold, n=30; *hyx* RNAi, 0.20-fold, n=34. *hyx* knockdown was driven by *insc*-Gal4 in A-B and G-H. NSCs are circled by white-dotted lines. Centrosomes are pointed by arrows. Scale bars: 5 μm.

**Figure S6.**
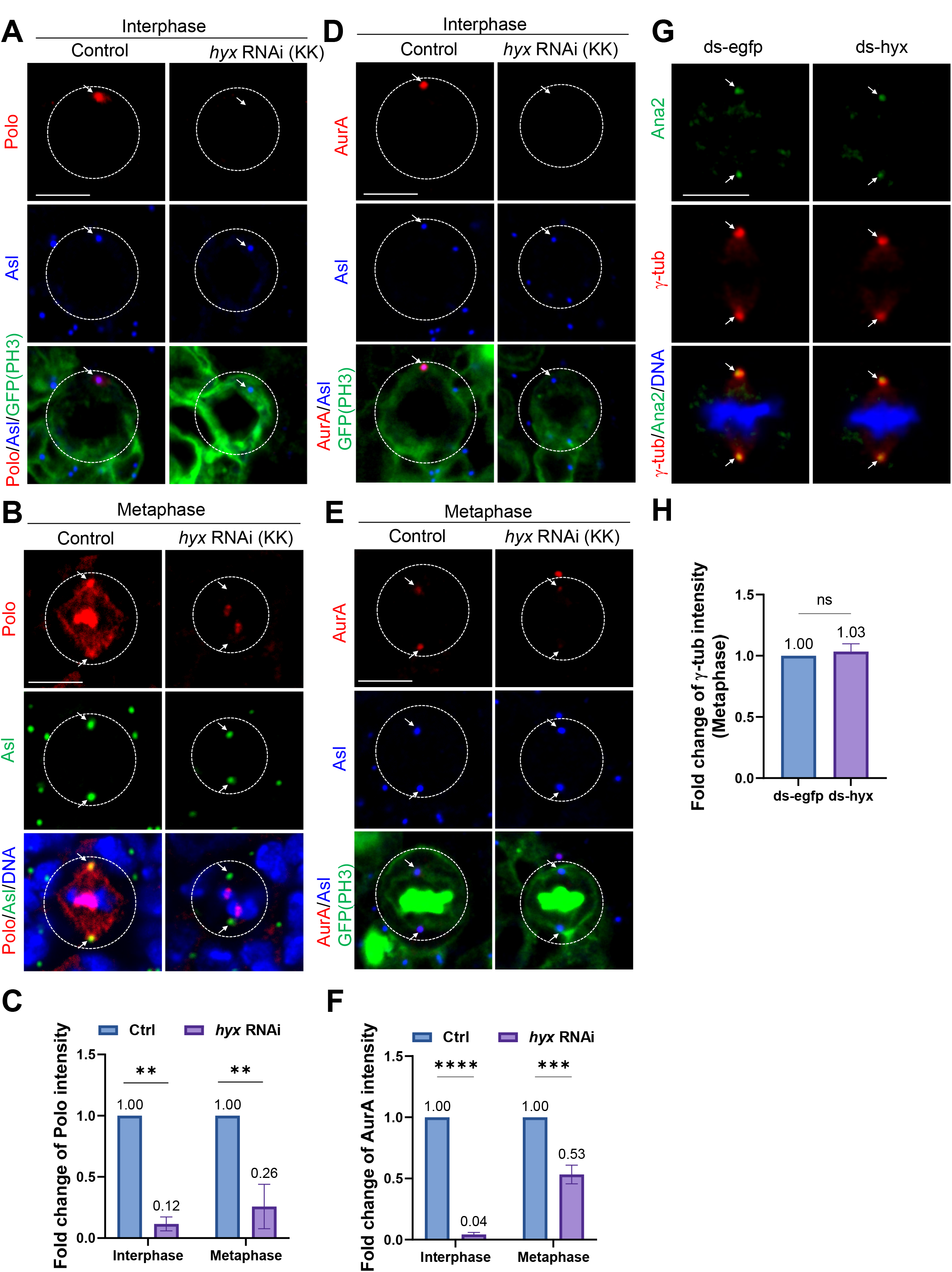
Polo and AurA were delocalized from the centrosomes upon *hyx* knockdown in NSCs and S2 cells. (A) Interphase NSCs of control (*UAS-β-Gal* RNAi; n=29) and *hyx* RNAi (KK/V103555; n=26) were labelled with Polo, Asl, and PH3. (B) Metaphase NSCs of control (*UAS-β-Gal* RNAi; Polo present at the centrosomes in 95.0% of NSCs, n=20) and *hyx* RNAi (KK/V103555; n=23) were labelled for Polo, Asl, and DNA. (C) Quantification graph showing the fold change of Polo intensity in A and B. Interphase: control, 1-fold, n=29; *hyx* RNAi, 0.12-fold, n=26. Metaphase: control, 1-fold, n=20; *hyx* RNAi, 0.26-fold, n=24. (D) Interphase NSCs of control (*UAS-β-Gal* RNAi; AurA present at the centrosomes in 96.6% of NSCs, n=29) and *hyx* RNAi (KK/V103555; n=24) were labelled for AurA, Asl, GFP, and PH3. (E) Metaphase NSCs of control (*UAS-β-Gal* RNAi; AurA observed at the centrosomes in 96.4% of NSCs, n=28) and *hyx* RNAi (KK/V103555; n=30) were labelled for AurA, Asl, GFP, and PH3. Polo and AurA are properly localized in all control NSCs in A-E. (F) Quantification graph showing the fold change of AurA intensity in D and E. Interphase: control, 1-fold, n=29; *hyx* RNAi, 0.04-fold, n=24. Metaphase: control, 1-fold, n=28; *hyx* RNAi, 0.53-fold, n=30. (G) Metaphase cells from ds-*egfp*-treated S2 cells and ds-*hyx*-treated S2 cells were labelled for Ana2, γ-tub, and DNA. (H) Quantification graph showing the fold change of Ana2 intensity in G. ds-*egfp*, 1-fold, n=38; ds-*hyx*, 0.83-fold, n=30. (I) Quantification graph showing the fold change of γ-tub intensity in G. ds-*egfp*, 1-fold, n=69; ds-*hyx*, 1.03-fold, n=52. *hyx* knockdown was under the control of *insc*-Gal4 in A-F. NSCs are outlined by white-dotted lines. Centrosomes are pointed by arrows. Scale bars: 5 μm.

## REFERENCES

1. Agarwal, S. K., W. F. Simonds and S. J. Marx (2008). “The parafibromin tumor suppressor protein interacts with actin-binding proteins actinin-2 and actinin-3.” Molecular cancer 7: 65–65.

2. Atwood, S. X., C. Chabu, R. R. Penkert, C. Q. Doe and K. E. Prehoda (2007). “Cdc42 acts downstream of Bazooka to regulate neuroblast polarity through Par-6 aPKC.” Journal of cell science 120(Pt 18): 3200–3206.

3. Bahrampour, S. and S. Thor (2016). “Ctr9, a Key Component of the Paf1 Complex, Affects Proliferation and Terminal Differentiation in the Developing Drosophila Nervous System.” G3 (Bethesda) 6(10): 3229–3239.

4. Bello, B., H. Reichert and F. Hirth (2006). “The brain tumor gene negatively regulates neural progenitor cell proliferation in the larval central brain of Drosophila.” Development 133(14): 2639–2648.

5. Bello, B. C., N. Izergina, E. Caussinus and H. Reichert (2008). “Amplification of neural stem cell proliferation by intermediate progenitor cells in Drosophila brain development.” Neural Develop 3: 5.

6. Berdnik, D. and J. A. Knoblich (2002). “Drosophila Aurora-A Is Required for Centrosome Maturation and Actin-Dependent Asymmetric Protein Localization during Mitosis.” Current Biology 12(8): 640–647.

7. Betschinger, J., K. Mechtler and J. A. Knoblich (2006). “Asymmetric Segregation of the Tumor Suppressor Brat Regulates Self-Renewal in Drosophila Neural Stem Cells.” Cell 124(6): 1241–1253.

8. Boone, J. Q. and C. Q. Doe (2008). “Identification of Drosophila type II neuroblast lineages containing transit amplifying ganglion mother cells.” Dev Neurobiol 68(9): 1185–1195.

9. Bowman, S. K., R. A. Neumüller, M. Novatchkova, Q. Du and J. A. Knoblich (2006). “The Drosophila NuMA Homolog Mud regulates spindle orientation in asymmetric cell division.” Dev Cell 10(6): 731–742.

10. Bowman, S. K., V. Rolland, J. Betschinger, K. A. Kinsey, G. Emery and J. A. Knoblich (2008). “The tumor suppressors Brat and Numb regulate transit-amplifying neuroblast lineages in Drosophila.” Dev Cell 14(4): 535–546.

11. Caussinus, E. and C. Gonzalez (2005). “Induction of tumor growth by altered stem-cell asymmetric division in Drosophila melanogaster.” Nat Genet 37(10): 1125–1129.

12. Chabu, C. and C. Q. Doe (2009). “Twins/PP2A regulates aPKC to control neuroblast cell polarity and self-renewal.” Developmental Biology 330(2): 399–405.

13. Chang, K. C., C. Wang and H. Wang (2012). “Balancing self-renewal and differentiation by asymmetric division: insights from brain tumor suppressors in Drosophila neural stem cells.” Bioessays 34(4): 301–310.

14. Chaturvedi, D., M. Inaba, S. Scoggin and M. Buszczak (2016). “Drosophila CG2469 Encodes a Homolog of Human CTR9 and Is Essential for Development.” G3 (Bethesda) 6(12): 3849–3857.

15. Chen, K., C. T. Koe, Z. B. Xing, X. Tian, F. Rossi, C. Wang, Q. Tang, W. Zong, W. J. Hong, R. Taneja, F. Yu, C. Gonzalez, C. Wu, S. Endow and H. Wang (2016). “Arl2- and Msps-dependent microtubule growth governs asymmetric division.” Journal of Cell Biology 212(6): 661–676.

16. Choksi, S. P., T. D. Southall, T. Bossing, K. Edoff, E. de Wit, B. E. Fischer, B. van Steensel, G. Micklem and A. H. Brand (2006). “Prospero acts as a binary switch between self-renewal and differentiation in Drosophila neural stem cells.” Dev Cell 11(6): 775–789.

17. Costa, P. J. and K. M. Arndt (2000). “Synthetic Lethal Interactions Suggest a Role for the Saccharomyces cerevisiae Rtf1 Protein in Transcription Elongation.” Genetics 156(2): 535–547.

18. Doe, C. Q. (2008). “Neural stem cells: balancing self-renewal with differentiation.” Development 135(9): 1575–1587.

19. Dzhindzhev, N. S., Q. D. Yu, K. Weiskopf, G. Tzolovsky, I. Cunha-Ferreira, M. Riparbelli, A. Rodrigues-Martins, M. Bettencourt-Dias, G. Callaini and D. M. Glover (2010). “Asterless is a scaffold for the onset of centriole assembly.” Nature 467(7316): 714–718.

20. Gartenmann, L., C. C. Vicente, A. Wainman, Z. A. Novak, B. Sieber, J. H. Richens and J. W. Raff (2020). “Drosophila Sas-6, Ana2 and Sas-4 self-organise into macromolecular structures that can be used to probe centriole and centrosome assembly.” Journal of Cell Science 133(12).

21. Giet, R., D. McLean, S. Descamps, M. J. Lee, J. W. Raff, C. Prigent and D. M. Glover (2002). “Drosophila Aurora A kinase is required to localize D-TACC to centrosomes and to regulate astral microtubules.” J Cell Biol 156(3): 437–451.

22. Gonzalez, C. (2007). “Spindle orientation, asymmetric division and tumour suppression in Drosophila stem cells.” Nat Rev Genet 8(6): 462–472.

23. Hannaford, M. R., A. Ramat, N. Loyer and J. Januschke (2018). “aPKC-mediated displacement and actomyosin-mediated retention polarize Miranda in Drosophila neuroblasts.” eLife 7: e29939.

24. Ikeshima-Kataoka, H., J. B. Skeath, Y.-i. Nabeshima, C. Q. Doe and F. Matsuzaki (1997). “Miranda directs Prospero to a daughter cell during Drosophila asymmetric divisions.” Nature 390(6660): 625–629.

25. Izumi, Y., N. Ohta, K. Hisata, T. Raabe and F. Matsuzaki (2006). “Drosophila Pins-binding protein Mud regulates spindle-polarity coupling and centrosome organization.” Nat Cell Biol 8(6): 586–593.

26. Januschke, J. and C. Gonzalez (2008). “Drosophila asymmetric division, polarity and cancer.” Oncogene 27(55): 6994–7002.

27. Januschke, J. and C. Gonzalez (2010). “The interphase microtubule aster is a determinant of asymmetric division orientation in Drosophila neuroblasts.” Journal of Cell Biology 188(5): 693–706.

28. Jo, J.-H., T.-M. Chung, H. Youn and J.-Y. Yoo (2014). “Cytoplasmic parafibromin/hCdc73 targets and destabilizes p53 mRNA to control p53-mediated apoptosis.” Nature Communications 5(1): 5433.

29. Kelsom, C. and W. Lu (2012). “Uncovering the link between malfunctions in Drosophila neuroblast asymmetric cell division and tumorigenesis.” Cell & Bioscience 2(1): 38.

30. Knoblich, J. A. (2010). “Asymmetric cell division: recent developments and their implications for tumour biology.” Nat Rev Mol Cell Biol 11(12): 849–860.

31. Koe, C. T. and H. Wang Asymmetric Cell Division in Drosophila Neuroblasts. eLS: 1–14.

32. Krahn, M. P., D. Egger-Adam and A. Wodarz (2009). “PP2A Antagonizes Phosphorylation of Bazooka by PAR-1 to Control Apical-Basal Polarity in Dividing Embryonic Neuroblasts.” Developmental Cell 16(6): 901–908.

33. Kraut, R., W. Chia, L. Y. Jan, Y. N. Jan and J. A. Knoblich (1996). “Role of inscuteable in orienting asymmetric cell divisions in Drosophila.” Nature 383(6595): 50–55.

34. Kubota, Y., K. Tsuyama, Y. Takabayashi, N. Haruta, R. Maruyama, N. Iida and A. Sugimoto (2014). “The PAF1 complex is involved in embryonic epidermal morphogenesis in Caenorhabditis elegans.” Developmental Biology 391(1): 43–53.

35. Lee, C.-Y., R. O. Andersen, C. Cabernard, L. Manning, K. D. Tran, M. J. Lanskey, A. Bashirullah and C. Q. Doe (2006). “Drosophila Aurora-A kinase inhibits neuroblast self-renewal by regulating aPKC/Numb cortical polarity and spindle orientation.” Genes & Development 20(24): 3464–3474.

36. Lee, C.-Y., K. J. Robinson and C. Q. Doe (2006). “Lgl, Pins and aPKC regulate neuroblast self-renewal versus differentiation.” Nature 439(7076): 594–598.

37. Lee, C.-Y., B. D. Wilkinson, S. E. Siegrist, R. P. Wharton and C. Q. Doe (2006). “Brat Is a Miranda Cargo Protein that Promotes Neuronal Differentiation and Inhibits Neuroblast Self-Renewal.” Developmental Cell 10(4): 441–449.

38. Lee, C. Y., R. O. Andersen, C. Cabernard, L. Manning, K. D. Tran, M. J. Lanskey, A. Bashirullah and C. Q. Doe (2006). “Drosophila Aurora-A kinase inhibits neuroblast self-renewal by regulating aPKC/Numb cortical polarity and spindle orientation.” Genes Dev 20(24): 3464–3474.

39. Lee, M. J., F. Gergely, K. Jeffers, S. Y. Peak-Chew and J. W. Raff (2001). “Msps/XMAP215 interacts with the centrosomal protein D-TACC to regulate microtubule behaviour.” Nature Cell Biology 3(7): 643–649.

40. Lee, T. and L. Luo (1999). “Mosaic analysis with a repressible cell marker for studies of gene function in neuronal morphogenesis.” Neuron 22(3): 451–461.

41. Li, S., C. T. Koe, S. T. Tay, A. L. K. Tan, S. Zhang, Y. Zhang, P. Tan, W.-K. Sung and H. Wang (2017). “An intrinsic mechanism controls reactivation of neural stem cells by spindle matrix proteins.” Nature Communications 8(1).

42. Lin, L., J. H. Zhang, L. M. Panicker and W. F. Simonds (2008). “The parafibromin tumor suppressor protein inhibits cell proliferation by repression of the c-myc proto-oncogene.” Proc Natl Acad Sci U S A 105(45): 17420–17425.

43. Lu, B., M. Rothenberg, L. Y. Jan and Y. N. Jan (1998). “Partner of Numb colocalizes with Numb during mitosis and directs Numb asymmetric localization in Drosophila neural and muscle progenitors.” Cell 95(2): 225–235.

44. Ly, P. T., Y. S. Tan, C. T. Koe, Y. Zhang, G. Xie, S. Endow, W. M. Deng, F. Yu and H. Wang (2019). “CRL4Mahj E3 ubiquitin ligase promotes neural stem cell reactivation.” PLoS Biol 17(6): e3000276.

45. Matsuzaki, F., T. Ohshiro, H. Ikeshima-Kataoka and H. Izumi (1998). “miranda localizes staufen and prospero asymmetrically in mitotic neuroblasts and epithelial cells in early Drosophila embryogenesis.” Development 125(20): 4089–4098.

46. Morin, X., R. Daneman, M. Zavortink and W. Chia (2001). “A protein trap strategy to detect GFP-tagged proteins expressed from their endogenous loci in Drosophila.” Proc Natl Acad Sci U S A 98(26): 15050–15055.

47. Moritz, M., M. B. Braunfeld, J. W. Sedat, B. Alberts and D. A. Agard (1995). “Microtubule nucleation by gamma-tubulin-containing rings in the centrosome.” Nature 378(6557): 638–640.

48. Mosimann, C., G. Hausmann and K. Basler (2006). “Parafibromin/Hyrax Activates Wnt/Wg Target Gene Transcription by Direct Association with β-catenin/Armadillo.” Cell 125(2): 327–341.

49. Mueller, C. L. and J. A. Jaehning (2002). “Ctr9, Rtf1, and Leo1 Are Components of the Paf1/RNA Polymerase II Complex.” Molecular and Cellular Biology 22(7): 1971.

50. Mueller, C. L., S. E. Porter, M. G. Hoffman and J. A. Jaehning (2004). “The Paf1 Complex Has Functions Independent of Actively Transcribing RNA Polymerase II.” Molecular Cell 14(4): 447–456.

51. Neumüller, R. A. and J. A. Knoblich (2009). “Dividing cellular asymmetry: asymmetric cell division and its implications for stem cells and cancer.” Genes & Development 23(23): 2675–2699.

52. Neumüller, R. A., C. Richter, A. Fischer, M. Novatchkova, K. G. Neumüller and J. A. Knoblich (2011). “Genome-wide analysis of self-renewal in Drosophila neural stem cells by transgenic RNAi.” Cell stem cell 8(5): 580–593.

53. Newey, P. J., M. R. Bowl and R. V. Thakker (2009). “Parafibromin--functional insights.” J Intern Med 266(1): 84–98.

54. Novak, Zsofia A., A. Wainman, L. Gartenmann and Jordan W. Raff (2016). “Cdk1 Phosphorylates Drosophila Sas-4 to Recruit Polo to Daughter Centrioles and Convert Them to Centrosomes.” Developmental Cell 37(6): 545–557.

55. Ogawa, H., N. Ohta, W. Moon and F. Matsuzaki (2009). “Protein phosphatase 2A negatively regulates aPKC signaling by modulating phosphorylation of Par-6 in Drosophila neuroblast asymmetric divisions.” J Cell Sci 122(Pt 18): 3242–3249.

56. Penheiter, K. L., T. M. Washburn, S. E. Porter, M. G. Hoffman and J. A. Jaehning (2005). “A Posttranscriptional Role for the Yeast Paf1-RNA Polymerase II Complex Is Revealed by Identification of Primary Targets.” Molecular Cell 20(2): 213–223.

57. Petronczki, M. and J. A. Knoblich (2001). “DmPAR-6 directs epithelial polarity and asymmetric cell division of neuroblasts in Drosophila.” Nature Cell Biology 3(1): 43–49.

58. Porzionato, A., V. Macchi, L. Barzon, G. Masi, M. Iacobone, A. Parenti, G. Palu and R. De Caro (2006). “Immunohistochemical assessment of parafibromin in mouse and human tissues.” J Anat 209(6): 817–827.

59. Porzionato, A., V. Macchi, L. Barzon, G. Masi, M. Iacobone, A. Parenti, G. Palù and R. De Caro (2006). “Immunohistochemical assessment of parafibromin in mouse and human tissues.” Journal of anatomy 209(6): 817–827.

60. Rea, S., F. Eisenhaber, D. O’Carroll, B. D. Strahl, Z. W. Sun, M. Schmid, S. Opravil, K. Mechtler, C. P. Ponting, C. D. Allis and T. Jenuwein (2000). “Regulation of chromatin structure by site-specific histone H3 methyltransferases.” Nature 406(6796): 593–599.

61. Roth, S. and R. Heintzmann (2016). “Optical photon reassignment with increased axial resolution by structured illumination.” Methods and Applications in Fluorescence 4(4):045005.

62. Rusan, N. M. and M. Peifer (2007). “A role for a novel centrosome cycle in asymmetric cell division.” The Journal of cell biology 177(1): 13–20.

63. Schaefer, M., M. Petronczki, D. Dorner, M. Forte and J. A. Knoblich (2001). “Heterotrimeric G Proteins Direct Two Modes of Asymmetric Cell Division in the Drosophila Nervous System.” Cell 107(2): 183–194.

64. Schober, M., M. Schaefer and J. A. Knoblich (1999). “Bazooka recruits Inscuteable to orient asymmetric cell divisions in Drosophila neuroblasts.” Nature 402(6761): 548–551.

65. Selvarajan, S., L. H. Sii, A. Lee, G. Yip, B. H. Bay, M. H. Tan, B. T. Teh and P. H. Tan (2008). “Parafibromin expression in breast cancer: a novel marker for prognostication?” J Clin Pathol 61(1): 64–67.

66. Shen, C.-P., L. Y. Jan and Y. N. Jan (1997). “Miranda Is Required for the Asymmetric Localization of Prospero during Mitosis in Drosophila.” Cell 90(3): 449–458.

67. Sunkel, C. E., R. Gomes, P. Sampaio, J. Perdigão and C. González (1995). “Gamma-tubulin is required for the structure and function of the microtubule organizing centre in Drosophila neuroblasts.” The EMBO journal 14(1): 28–36.

68. Terada, Y., Y. Uetake and R. Kuriyama (2003). “Interaction of Aurora-A and centrosomin at the microtubule-nucleating site in Drosophila and mammalian cells.” The Journal of cell biology 162(5): 757–763.

69. Van Oss, S. B., C. E. Cucinotta and K. M. Arndt (2017). “Emerging Insights into the Roles of the Paf1 Complex in Gene Regulation.” Trends Biochem Sci 42(10): 788–798.

70. Varmark, H., S. Llamazares, E. Rebollo, B. Lange, J. Reina, H. Schwarz and C. Gonzalez (2007). “Asterless Is a Centriolar Protein Required for Centrosome Function and Embryo Development in Drosophila.” Current Biology 17(20): 1735–1745.

71. Wang, C., K. C. Chang, G. Somers, D. Virshup, B. T. Ang, C. Tang, F. Yu and H. Wang (2009). “Protein phosphatase 2A regulates self-renewal of Drosophilaneural stem cells.” Development 136(13): 2287–2296.

72. Wang, C., S. Li, J. Januschke, F. Rossi, Y. Izumi, G. Garcia-Alvarez, S. S. Gwee, S. B. Soon, H. K. Sidhu, F. Yu, F. Matsuzaki, C. Gonzalez and H. Wang (2011). “An ana2/ctp/mud complex regulates spindle orientation in Drosophila neuroblasts.” Dev Cell 21(3): 520–533.

73. Wang, C., S. Li, J. Januschke, F. Rossi, Y. Izumi, G. Garcia-Alvarez, Serene Sze L. Gwee, Swee B. Soon, Harpreet K. Sidhu, F. Yu, F. Matsuzaki, C. Gonzalez and H. Wang (2011). “An Ana2/Ctp/Mud Complex Regulates Spindle Orientation in Drosophila Neuroblasts.” Developmental Cell 21(3): 520–533.

74. Wang, G., Q. Jiang and C. Zhang (2014). “The role of mitotic kinases in coupling the centrosome cycle with the assembly of the mitotic spindle.” J Cell Sci 127(Pt 19): 4111–4122.

75. Wang, H., Y. Ouyang, W. G. Somers, W. Chia and B. Lu (2007). “Polo inhibits progenitor self-renewal and regulates Numb asymmetry by phosphorylating Pon.” Nature 449(7158): 96–100.

76. Wang, H., G. W. Somers, A. Bashirullah, U. Heberlein, F. Yu and W. Chia (2006). “Aurora-A acts as a tumor suppressor and regulates self-renewal of Drosophila neuroblasts.” Genes & Development 20(24): 3453–3463.

77. Wang, H., G. W. Somers, A. Bashirullah, U. Heberlein, F. Yu and W. Chia (2006). “Aurora-A acts as a tumor suppressor and regulates self-renewal of Drosophila neuroblasts.” Genes Dev 20(24): 3453–3463.

78. Wang, P., M. R. Bowl, S. Bender, J. Peng, L. Farber, J. Chen, A. Ali, Z. Zhang, A. S. Alberts, R. V. Thakker, A. Shilatifard, B. O. Williams and B. T. Teh (2008). “Parafibromin, a component of the human PAF complex, regulates growth factors and is required for embryonic development and survival in adult mice.” Molecular and cellular biology 28(9): 2930–2940.

79. Wei, Y., L. Yu, J. Bowen, M. A. Gorovsky and C. D. Allis (1999). “Phosphorylation of histone H3 is required for proper chromosome condensation and segregation.” Cell 97(1): 99–109.

80. Weng, M., K. L. Golden and C. Y. Lee (2010). “dFezf/Earmuff maintains the restricted developmental potential of intermediate neural progenitors in Drosophila.” Dev Cell 18(1): 126–135.

81. Wodarz, A., A. Ramrath, A. Grimm and E. Knust (2000). “Drosophila atypical protein kinase C associates with Bazooka and controls polarity of epithelia and neuroblasts.” The Journal of cell biology 150(6): 1361–1374.

82. Wodarz, A., A. Ramrath, U. Kuchinke and E. Knust (1999). “Bazooka provides an apical cue for Inscuteable localization in Drosophila neuroblasts.” Nature 402(6761): 544–547.

83. Woodard, G. E., L. Lin, J. H. Zhang, S. K. Agarwal, S. J. Marx and W. F. Simonds (2005). “Parafibromin, product of the hyperparathyroidism-jaw tumor syndrome gene HRPT2, regulates cyclin D1/PRAD1 expression.” Oncogene 24(7): 1272–1276.

84. Wu, P.-S., B. Egger and A. H. Brand (2008). “Asymmetric stem cell division: Lessons from Drosophila.” Seminars in Cell & Developmental Biology 19(3): 283–293.

85. Yang, Y.-J., J.-W. Han, H.-D. Youn and E.-J. Cho (2010). “The tumor suppressor, parafibromin, mediates histone H3 K9 methylation for cyclin D1 repression.” Nucleic acids research 38(2): 382–390.

86. Yu, F., X. Morin, Y. Cai, X. Yang and W. Chia (2000). “Analysis of partner of inscuteable, a Novel Player of Drosophila Asymmetric Divisions, Reveals Two Distinct Steps in Inscuteable Apical Localization.” Cell 100(4): 399–409.

87. Yu, F., H. Wang, H. Qian, R. Kaushik, M. Bownes, X. Yang and W. Chia (2005). “Locomotion defects, together with Pins, regulates heterotrimeric G-protein signaling during Drosophila neuroblast asymmetric divisions.” Genes & development 19(11): 1341–1353.

88. Zhang, Z., X. F. Yang, K. Q. Huang, L. Ren, W. F. Gou, D. F. Shen, S. Zhao, H. Z. Sun, Y. Takano and H. C. Zheng (2015). “The clinicopathological significances and biological functions of parafibromin expression in head and neck squamous cell carcinomas.” Tumour Biol 36(12): 9487–9497.

89. Zheng, H. C., H. Takahashi, X. H. Li, T. Hara, S. Masuda, Y. F. Guan and Y. Takano (2008). “Downregulated parafibromin expression is a promising marker for pathogenesis, invasion, metastasis and prognosis of gastric carcinomas.” Virchows Arch 452(2): 147–155.

90. Zhu, B., S. S. Mandal, A.-D. Pham, Y. Zheng, H. Erdjument-Bromage, S. K. Batra, P. Tempst and D. Reinberg (2005). “The human PAF complex coordinates transcription with events downstream of RNA synthesis.” Genes & Development 19(14): 1668–1673.

